# Heterochronic scaling of neurogenesis for species-specific dosing of cortical excitatory subtypes

**DOI:** 10.1101/2025.04.10.648214

**Authors:** Yuki Y. Yamauchi, Xuanhao D. Sheu, Rafat Tarfder, Takuma Kumamoto, Jun Hatakeyama, Haruka Sato, Pauline Rouillard, Merve Bilgic, Shuto Deguchi, Tomonori Nakamura, Yusuke Kishi, Kazuo Emoto, Ikuo K. Suzuki

## Abstract

Mammals share a laminar cerebral cortex, with excitatory neuron subtypes organized in distinct layers. Although this framework is conserved, subtype balance varies markedly between species due to unknown mechanisms. This study shows that species-specific neuronal composition arises from non-uniform scaling of the temporal dynamics of neurogenesis. Comparative histology of eight mammalian species revealed a significant, rat-specific expansion of the cortical deeper layer (DL). This species difference results from a specific extension of the early neurogenetic phase for DL neuron production before transitioning to the upper layer (UL) in rats, as confirmed by birthdating and single-cell transcriptomics. The duration of DL neuron production is regulated by a genetic program controlling progenitor aging, including canonical Wnt signaling. Comparative single-cell transcriptomics revealed that cortical progenitors in rats exhibit significantly elevated Wnt ligand expression. Thus, while sequential cortical neurogenesis is conserved, its progression is non-uniformly scaled in each species. Such precise heterochronic fine-tuning allows evolutionary refinement of cellular configuration without drastic remodeling of the conserved corticogenesis program.

**Teaser:** Differential temporal progression of neural progenitor program contributes to species-specific cellular composition of mammalian cerebral cortex

## Introduction

Animals adapt their neural circuits to enhance survival in specific ecological niches(*1*, *2*). In mammals, a highly developed cerebral cortex underlies a wide range of brain functions and ultimately leads to advanced cognitive abilities, especially in humans. The mammalian cortex is composed of a finely balanced repertoire of neuronal subtypes, organized into six distinct tangential layers(*3–5*).

These include excitatory subtypes specific to certain cortical layers: deeper layers (DL; layers 5 and 6) are predominantly occupied by extra-cortical projection neurons, layer 4 by input-recipient neurons, and layers 2/3 by intra-cortical projection neurons. The diversity and laminar distribution of excitatory neuron subtypes are generally conserved across mammalian species, but significant quantitative variations do occur(*5–7*), presumably relevant to the functional adaptations.

The mechanisms regulating neuronal dosage and subtype balance remain poorly understood. Cortical excitatory neurons are generated by radial glial (RG) progenitors, also known as apical progenitors, in the ventricular zone (VZ)(*8–10*) directly or indirectly through intermediate progenitors. These neurons are produced in a stereotyped temporal sequence, with deeper-layer neurons forming earlier than more superficial layer neurons(*11–14*). This conserved neurogenic sequence has undergone significant quantitative modifications in certain species or animal phyla. For instance, the human cortex exhibits an expansion of upper layer (UL; layers 2–4) neurons compared to other species(*7*, *11*, *15–19*). This increase is linked to an extremely extended late neurogenic phase for UL neuron production. Whether similar temporal adaptations underlie other changes in cortical subtype composition during evolution remains unclear. Additionally, the specific aspects of cortical neurogenesis that are evolutionarily adaptable are yet to be determined.

Most studies of cortical evolution have concentrated on human-specific traits(*11*, *17*, *18*, *20*, *21*), whereas other mammalian lineages remain comparatively underexplored. Rodents, one of the most diverse mammalian orders, exhibit substantial variation in morphology, behavior, and brain organization(*22*, *23*). Among them, mice and rats—two of the most widely used experimental models—differ markedly in brain structure, volume, and behavior. These species diverged approximately 13.1 million years ago(*24*), and have since accumulated distinct evolutionary adaptations. Despite their central role in biomedical research, relatively few studies have addressed the evolutionary divergence of their cortical architecture(*25*).

In this study, we began by comparing cortical layer organization across eight mammalian species and uncovered an unexpected trend: rats possess an exceptionally thick DL, even relative to other rodents. To investigate the developmental basis of this species-specific neuronal composition, we analyzed excitatory cortical neurogenesis in rats and mice at single-cell resolution. Clonal lineage tracing demonstrated that rat RG progenitors generate a disproportionately higher number of DL neurons, while the number of UL neurons remains conserved. Complementary neuronal birthdating and single-cell RNA sequencing (scRNA-seq) revealed that temporal regulation of neurogenesis by RG progenitors scales in a non-uniform manner between species, ultimately producing distinct neuronal ensembles.

## Results

### DL-biased adaptation in the rodent cerebral cortex

To survey evolutionary variation in mammalian neocortical architecture, we histologically compared adult somatosensory cortices from eight mammalian species (**Figure 1A**, **Figure S1**). A general pattern emerged: gyrencephalic species exhibited proportionally thicker UL, whereas lissencephalic species, namely rodents, displayed thicker DL (**Figure 1A**, **Figure S1**), as reported previously(*26*). Within rodent species, there is a considerable variation in UL-DL ratio, with a most notable exception in rats, which have a significant bias of DL thickening. This contrasts with the human lineage, where UL thickness is widely attributed to enhanced UL production during the late neurogenetic phase. Despite intense interest in human cortical evolution, specific changes in other clades remain comparatively underexplored. DL expansion in rodents, most drastically observed in rats, thus provides a tractable model for evolutionary adaptation of cortical structure.

**Fig. 1.**
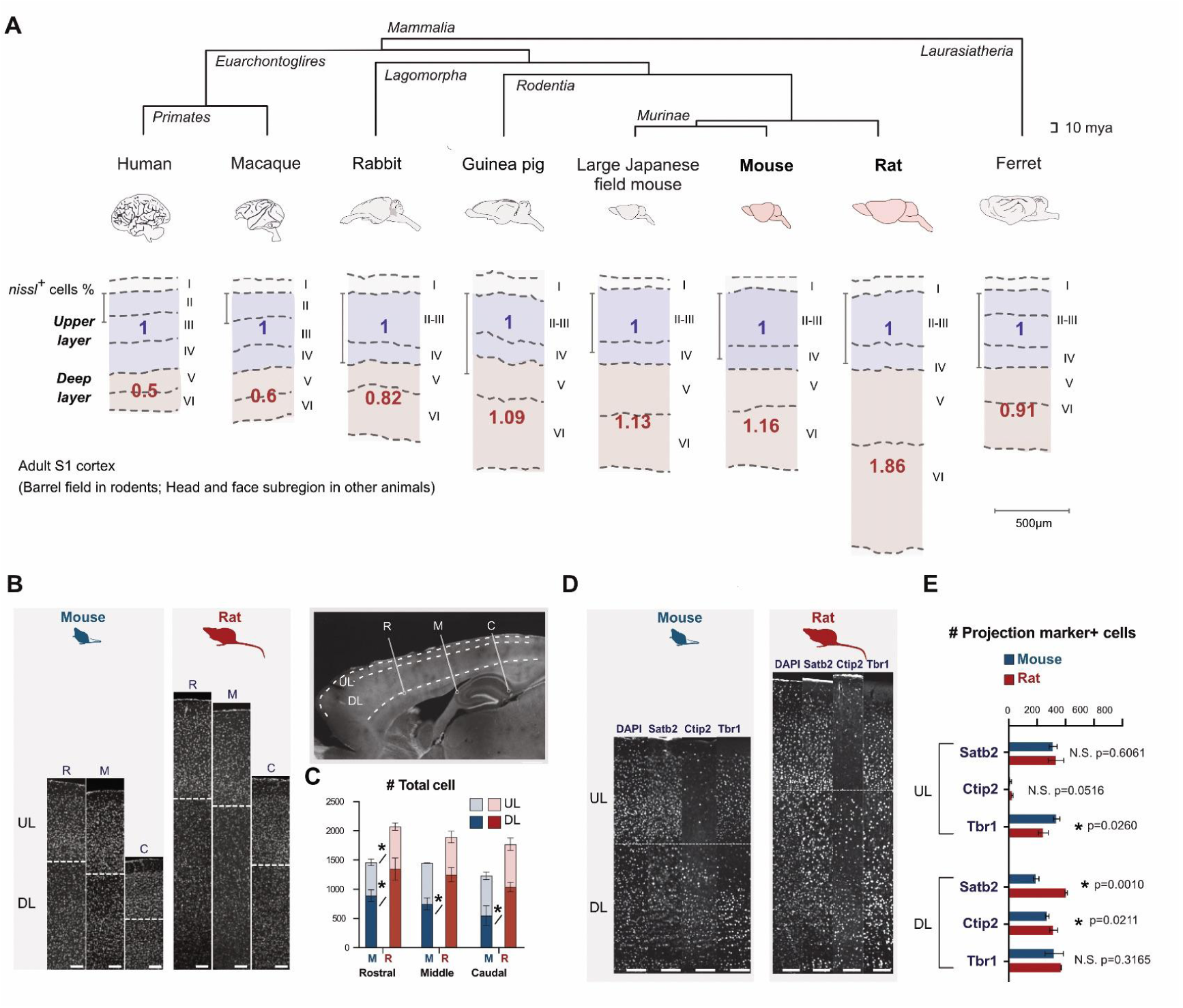
Histological comparison of the mammalian cerebral cortex highlighting the exceptional expansion of DL in rats. **A.** Phylogenetic tree of eight mammalian species with the upper-to-deep layer neuron ratio in the somatosensory cortex. Neuronal numbers were quantified using Nissl-stained brain sections from the corresponding cortical region at the adult stage. UL thickness was normalized across species to facilitate comparison of evolutionary variations in DL expansion. **B.** DAPI nuclear staining of the adult mouse and rat cerebral cortex. Three rostral-to-caudal planes are defined relative to the hippocampus. White dotted lines delineate the boundary between UL and DL. **C.** Total number of DAPI-labeled nuclei in the UL and DL across the three rostral-to-caudal levels in both species. Statistical significance (p < 0.05) is indicated by asterisks. **D.** Immunohistochemical detection of projection neuron-specific markers (Satb2, Ctip2, and Tbr1). White dotted lines delineate the boundary between UL and DL. **E.** Quantification of Satb2-, Ctip2-, and Tbr1-expressing neurons in the UL and DL of the mouse and rat primary somatosensory cortex (S1) at the adult stage. Data are presented as mean ± SEM (n = 3 samples per group); p values were calculated using Student’s t-test. Scale bars: 500 μm(A), 100 μm (B and D).

To probe these species differences, we performed detailed histology in mice and rats. Nuclear staining showed that overall laminar organization is conserved; however, rats possess a markedly thicker DL, which is less cell-dense than the UL, across rostral–caudal cortical regions (**Figure 1B**, **Figure S2**). Quantification confirmed that rats harbor more DL neurons than mice, whereas UL neuron numbers are relatively conserved (**Figure 1C**). We corroborated these findings by immunohistochemistry for the markers of excitatory subtypes with different axon projections: Satb2 (callosal projection neurons present in UL and a subset of DL; CPN), Ctip2 (corticospinal projection neurons, predominantly DL; CSPN), and Tbr1 (corticothalamic projection neurons, mainly DL; CThPN). In the primary somatosensory cortex, delineated by the characteristic barrel structures (**Figure S3**), rats displayed significantly higher numbers of Satb2⁺ and Ctip2⁺ DL neurons than mice, while counts of UL subtypes were nearly identical between species or higher in mice (**Figure 1D, E**). These DL-biased differences were evident already at an early postnatal stage P7 (**Figure S2**). Consistent with the expectation that DL-derived descending pathways would show larger species divergence than intracortical (primarily UL-derived) fibers, the corticospinal tract exhibited a substantially greater cross-sectional difference between mice and rats than did the corpus callosum (**Figure S2**). Together, these results indicate that, despite conserved laminar distribution, species divergence between mice and rats is characterized chiefly by an expansion of DL neuron populations.

### Individual cortical RG progenitors generate species-specific number of DL neurons

To investigate the developmental basis of species differences in cortical neuron balance, we studied the neurogenetic process in the two species. Since excitatory neurons are locally supplied by RG progenitors within the same tangential position, the clonally-related sister neurons are radially clustered over the multiple layers in a columnar space in the neocortex. Previous studies using Mosaic analysis with double markers (MADM) have shown that individual RG progenitors in mice generate an average of 8–9 clonal neurons, with 40–50% belonging to DL subtypes(*27*, *28*). To quantitatively compare the neuronal subtype production of individual progenitors in mice and rats in an equivalent manner, we used "hemi-lineage" labeling with retroviruses expressing fluorescent reporters(*28*) (See details for hemi-lineage labeling in **Figure S4A-B**). This method integrates a single copy of the retroviral vector into the host genome, permanently labeling progeny cells derived from one daughter cell of the initially infected host cell. The equally optimized titer of retroviruses expressing GFP and RFP were injected at the embryonic stage, with animals sacrificed at postnatal day 7 (P7) (**Figure 2A-B, Figure S4B**), when cortical neurogenesis is complete in both species (**Figure S4C**). GFP- and RFP-positive cells were sparsely distributed across cortical layers without overlap between clones, confirming spatial separation of labeled clones (**Figure S4B**).

**Figure 2.**
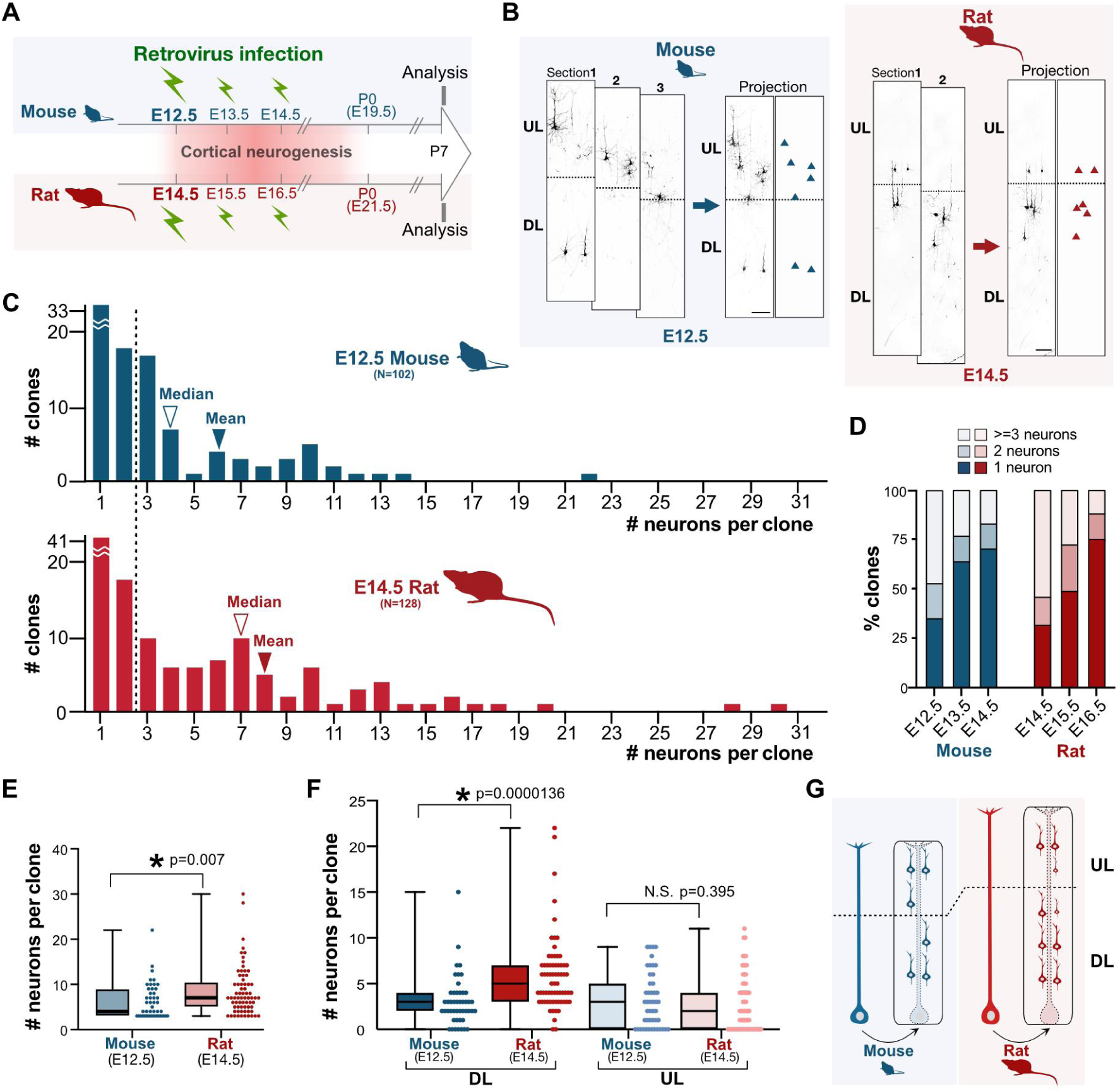
Clonal production of cortical excitatory neurons in mice and rats. **A**. Time course of clonal analysis in the mouse and rat cerebral cortices. Retroviral injections were performed at the onset of neurogenesis (E12.5 in mice and E14.5 in rats), and animals were sacrificed at P7 for analysis. **B**. Representative examples of cortical excitatory neuron clones in mice and rats. Scale bars: 100 μm. **C**. Distribution of clone sizes when retroviruses were injected at the onset of neurogenesis. Clonal sizes represent the number of neurons generated by individual RG progenitors. Median and mean of clone size are indicared by open and filled arrowheads, respectively. **D**. Percentage of clones containing one, two, or three or more neurons in mice and rats. **E**. Comparison of clone size between species. Only clones with three or more neurons were analyzed. **F**. Distributions of upper layer (UL) and deep layer (DL) neurons per clone. Rats exhibit a marked increase in the number of DL neurons per clone compared to mice, while UL neuron numbers remain similar between the species. **G**. Schematic diagram summarizing the clonal analysis results (mean number of neurons per clone), highlighting species-specific differences in DL neuron production. Statistical analysis: p-values calculated using Student’s t-test.

To compare whole neurogenetic processes between the two species, we examined the transition timing of RG progenitors from proliferative to neurogenic states; *i.e.* the onset of neuron production. At any given stage of development, RG progenitors are heterogeneous, comprising both proliferative and neurogenic subpopulations in specific ratios. The progenitor population gradually shifts from predominantly proliferative to a neurogenic mode. In mice, previous studies have shown that most cortical RG progenitors complete this transition between E11.5 and E12.5(*27*, *28*). To identify the corresponding stage in rats, we quantified the percentage of clones containing one or two neurons, as these indicate neurogenic progenitors at the time of retroviral injection (**Figure S4A**).

Retroviral infection at E12.5 in mice revealed that nearly half of the clones contained one or two neurons, consistent with earlier findings(*28*) (**Figure 2C** and **D**). A highly comparable pattern was observed in rats when the injection was performed at E14.5 (**Figure 2D**). In rats, neurogenic division became dominant at E15.5–E16.5, with over 50% of clones containing one or two neurons during these stages (**Figure 2D**). These results indicate that E12.5 in mice corresponds to E14.5 in rats as the onset of neurogenesis for most cortical RG progenitors. This correspondence is further supported by conserved neuron-to-progenitor ratios identified in immunohistological analyses (**Figure S4D**).

Next, we compared neuron production by individual progenitors from the onset of neurogenesis (E12.5 in mice, E14.5 in rats) to P7 (**Figure 2B-C**). Only clones comprising three or more neurons were considered, as smaller clones likely represent incomplete lineages (**Figure S2A**). In mice, the mean clonal size was 6.22 ± 3.88 (median 4), consistent with previous studies(*28*) (**Figure 2C** and **E**). In contrast, clonal size distribution showed a rightward skew in rats, with a mean of 8.05 ± 5.44 (median 7) (**Figure 2C** and **E**). Neurons were categorized into DL or UL subtypes based on somal position. Rats exhibited significantly larger numbers of DL neurons per clone compared to mice (5.90 ± 4.01 in rats vs. 3.04 ± 2.69 in mice), while UL neuron numbers were similar (2.62 ± 2.98 in rats vs. 3.11 ± 2.95 in mice) (**Figure 2F**). These results indicate that both species possess multipotent cortical RG progenitors capable of generating both DL and UL subtypes. However, neurogenic activity of each individual progenitor, particularly for DL subtype generation, is greater in rats than in mice (**Figure 2G**).

### Species-specific timeline of cortical neurogenesis; prolonged early neurogenic phase for DL subtypes in rats than in mice

We sought to understand the mechanism underlying such a species-specific dosage of DL neuron production. An important clue was obtained by clonal analysis at different time points of retroviral injection (**Figure 2A**, **Figure 3** and **Figure S5**). Retroviruses were injected at the onset of neurogenesis ("Day 1") and 24 hours ("Day 2") and 48 hours ("Day 3") later. Progeny cells were analyzed at P7, as in the other analyses (**Figure 3A**). The overall pattern of neurogenesis was consistent in both species. The proportion of monopotent clones containing only UL neurons increased progressively from Day 1 to Day 3, reflecting the inside-out sequence of cortical neurogenesis(*12*, *29*). However, a clear species-specific difference was observed: mice displayed a higher percentage of clones containing only UL neurons in injections on Days 2 and 3, while rats retained a greater proportion of multipotent clones capable of generating DL neurons, even with injections on Days 2 and 3 (**Figure 3A**). This difference was further supported by the mean number of DL neurons per clone. In rats, DL neurons were still observed when retroviruses were injected on Day 3, whereas in mice, DL neurons were absent in the corresponding condition (**Figure 3A and B**).

**Figure 3.**
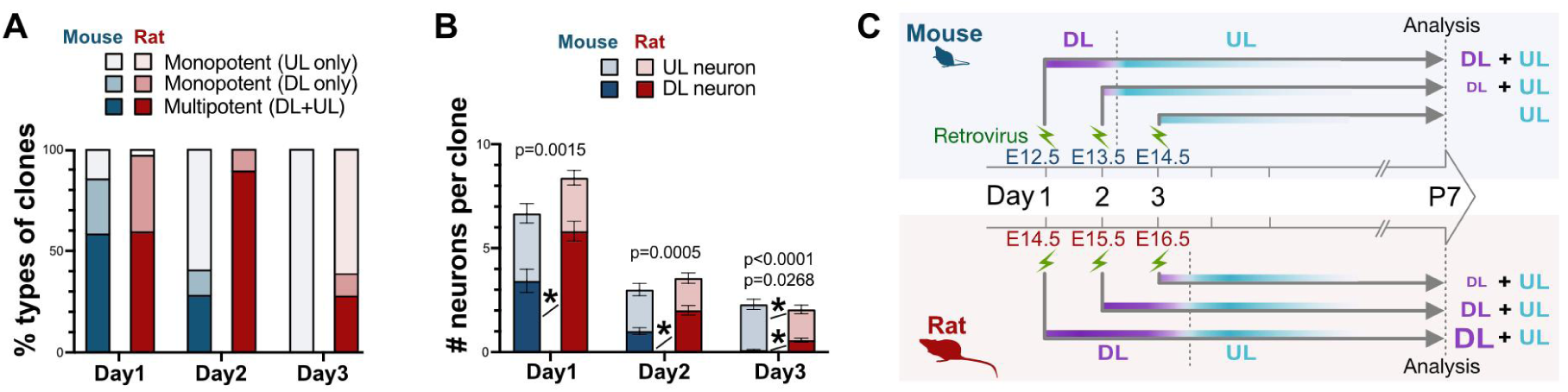
Temporal progression of cortical neurogenesis revealed by retroviral labeling of RG progenitors at distinct injection timings. **A**. Clones were categorized into three types: multipotent (DL + UL), monopotent (DL only), and monopotent (UL only). By Day 3 of neurogenesis, all mouse clones were UL-only monopotent type, whereas a significant proportion of rat clones remained multipotent when retroviruses were injected at the corresponding stage. **B.** Number of DL and UL neurons per clone after retrovirus injection at Days 1, 2, and 3 of neurogenesis in mice and rats. Rat clones contain DL neurons even at Day 3, unlike mouse clones, which transitioned entirely to UL neurons. Data are presented as mean ± SEM. Statistical significance was calculated using Student’s t-test. **C**. Summary illustration of clonal analysis with retroviral injections at different time points. Vertical dotted lines indicate the expected timing of the DL-to-UL subtype transition for the majority of RG progenitors in mice and rats.

These findings suggest that cortical neurogenesis follows distinct temporal programs in the two species. In rats, the early neurogenic phase, during which DL neurons are produced, is extended beyond Day 3. In contrast, mice complete the transition from DL to UL neuron production by Day 3 (**Figure 3C**).

To directly examine the timing of cortical neuron subtype generation in mice and rats, we conducted a birthdating analysis using the nucleotide analog 5-ethynyl-2’-deoxyuridine (EdU) (**Figure 4A**). EdU was administered to label neurons undergoing their final mitosis at specific developmental stages, from the onset of neurogenesis (Day 1; E12.5 in mice and E14.5 in rats) until one day before delivery (Day 7; E18.5 in mice and E20.5 in rats). Labeled neurons were subsequently analyzed for layer positions and marker expression at P7. The analysis revealed a significant species-specific difference in neuron generation timing based on the layer positions of EdU-labeled neurons. In mice, DL neurons were generated within a two-day window (Days 1–2) before transitioning to UL neuron production. In contrast, DL neurons in rats were produced over a prolonged four-day period (Days 1–4) (**Figure 4B-D**). The duration of UL neurogenesis was comparable between species, lasting four days before terminating during postnatal stages. This finding was supported by the quantification of subtype-specific markers (**Figure 4E-H** and **Figure S6**). These results indicate that the conserved inside-out temporal program of cortical neurogenesis is finetuned specifically in the early neurogenetic phase in a species-specific manner. This non-uniform finetuning is characterized by the extended duration of DL neuron production in rats before the transition to UL neuron production (**Figure 4I**).

**Figure 4.**
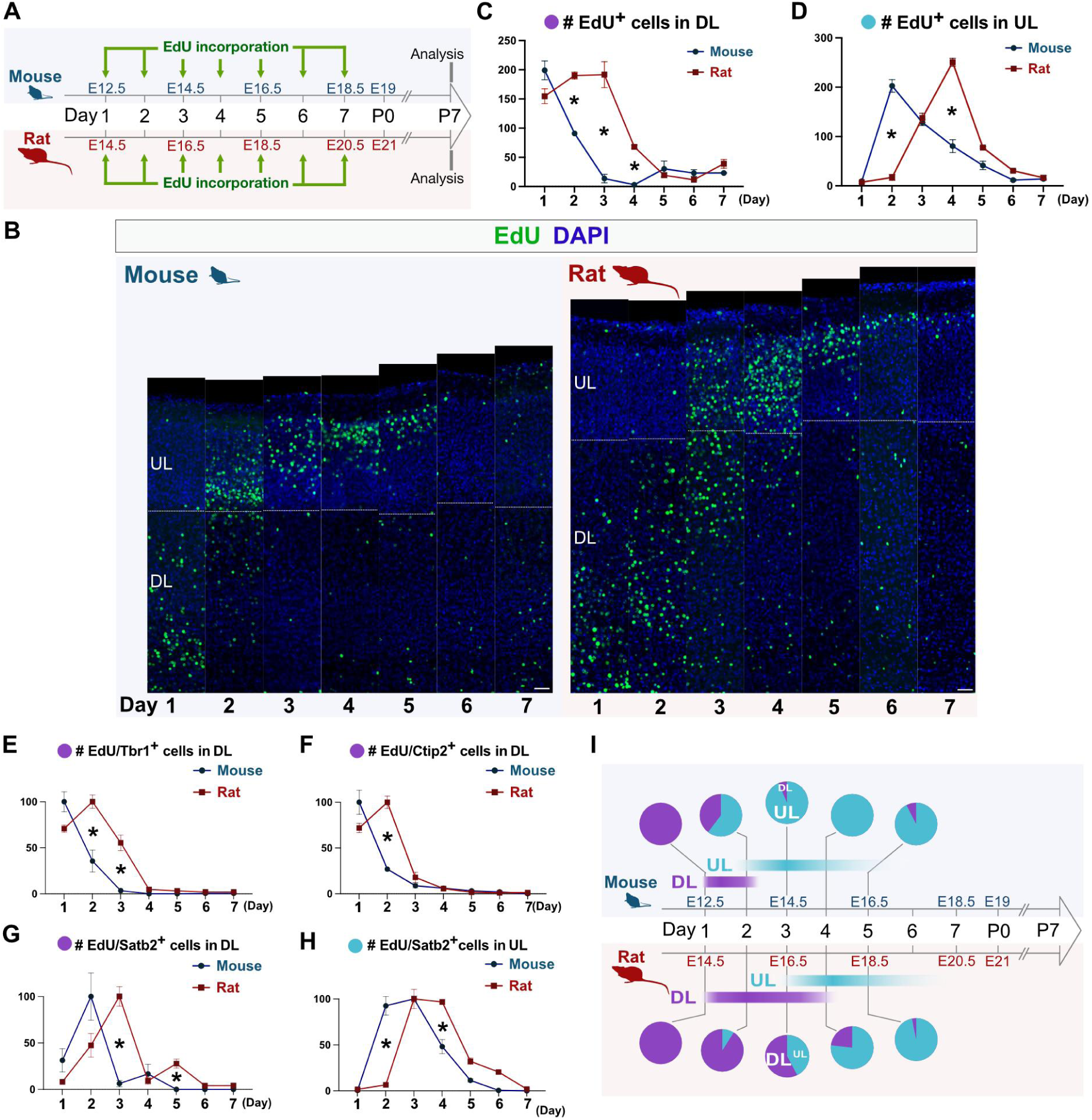
Birthdating of cortical excitatory neurons in mice and rats. **A**. Experimental design for birthdating analysis in mice and rats, showing different timing of EdU injections from the onset until the end of neurogenesis. **B**. Representative images of cortical columns in S1 with EdU-labeled neurons at the specified developmental stages. Scale bars: 50 μm. **C, D**. Quantification of DL (**C**) and UL subtypes (**D**) labeled with EdU at different stages. Asterisks indicate statistical significance (p < 0.05). **E-H**. Normalized counts of neurons colabeled for EdU and projection-specific markers. (**E**) Tbr1-positive DL neurons, (**F**) Ctip2-positive DL neurons, (**G**) Satb2-positive neurons located in DL, and (**H**) Satb2-positive neurons located in UL. **I**. Summary of neuronal birthdating in mice and rats, highlighting species-specific differences in the timing of DL and UL neuron production. Statistical analysis was performed using Welch’s t-test, with detailed results available in **Table S2**.

### Species-specific timing of DL to UL subtype transition confirmed by axon projection analysis

To further confirm the species-specific timing of the DL to UL subtype switch, characterized by somal location and marker expression (**Figures 3 and 4**), we examined the axon projection patterns of neurons generated during the mid-neurogenic phase (Days 2.5 and 3), since axonal projections are the defining feature of these subtypes (**Figure S7B**). We analyzed axon projections of the neurons generated at a certain timing by *in utero* electroporation (IUE) of a plasmid encoding the membrane-anchored fluorescent protein Achilles. This method labeled excitatory neurons in the primary somatosensory cortex and enabled axonal trajectory tracing at P7 (**Figure S7A**). The laminar distribution of labeled neurons electroporated at Day 2.5 closely matched the results of EdU pulse-labeling at Day 3, which demonstrated the greatest species-specific differences in neuronal composition (**Figure S7C**). IUE efficiently labeled axons for visualization in whole brains using tissue clearing (**Figure S7D**) and in slices with immunostaining of Achilles protein with anti-GFP antibody (**Figure S7E**). In mice, axons of neurons labeled at the mid-neurogenic stage predominantly projected to the corpus callosum. In contrast, in rats, axons from neurons labeled at the same stage projected to both intra- and extra-cortical targets, including the corpus callosum, thalamus, and descending brainstem structures such as the ipsilateral pyramidal tract (**Figure S7D–F**). Consistent with previous studies(*4*), neurons generated at Days 1–1.5 in mice projected to brainstem targets, indicating that the DL to UL subtype transition occurs between Days 1.5 and 2 in mice (**Figure S7E**).

These findings align with the birthdating results (**Figures 3 and 4**), where neurons born at the mid-neurogenic stage (Day 3) in mice were predominantly Satb2-positive UL subtypes in layer 2/3 (CPN). Conversely, in rats, neurons generated at the same stage included more Tbr1-positive CThPN and Ctip2-positive CSPN in layers 5–6. These observations confirm that the mid-neurogenic phase produces distinct neuronal subtypes in mice and rats, defined by axon projection patterns. The differences in axonal targeting reflect the species-specific timing of the DL to UL subtype switch (**Figure S7F**).

### Conservation in neural progenitor cycling and indirect neurogenesis in mice and rats

The species-specific difference in DL neuron production could theoretically arise from transient amplification of neuronal output through either cell cycle acceleration or an increased contribution of intermediate progenitors (IP) via indirect neurogenesis(*30–33*) during the early DL-producing phase. To explore these possibilities, we analyzed the density of mitotic progenitors in the VZ and subventricular zone (SVZ) of the future primary somatosensory field (**Figure 5A-C**). The temporal patterns of mitotic density in both the VZ and SVZ were strikingly conserved between the two species. Additionally, we assessed progenitors in the G2/S phase by administering EdU one hour before sacrifice (**Figure 5D** and **E**). No significant differences were observed in the density of G2/S-phase cells throughout the neurogenic period, except at the end of neurogenesis on Day 7. These findings suggest that cell cycle kinetics, as partly indicated by proportions of progenitors in the M and G2/S phases, are comparable between the two species, consistent with previous studies(*34*, *35*).

**Figure 5.**
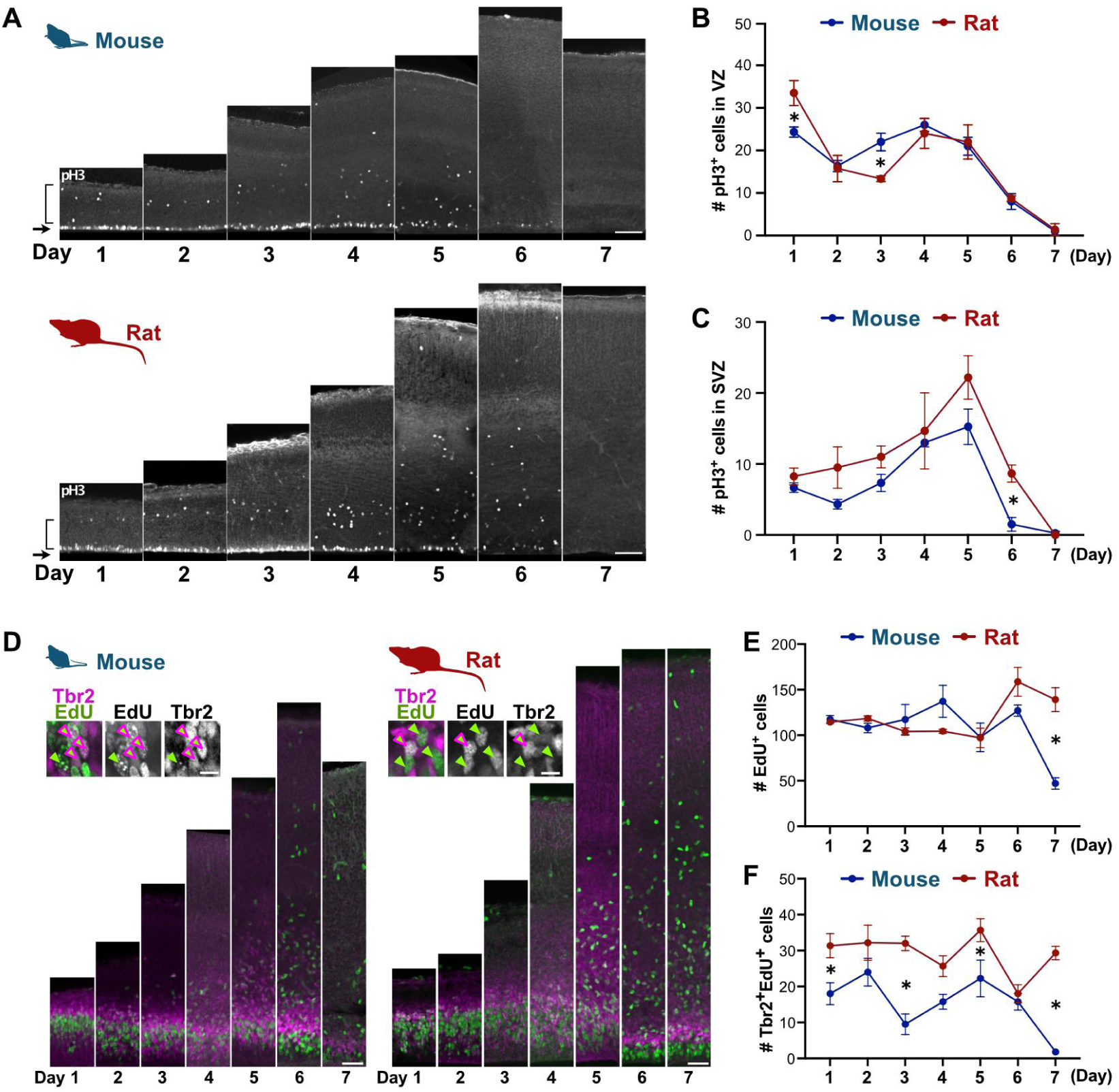
Dynamics of progenitor cell cycle and indirect neurogenesis activity in mouse and rat cortical neurogenesis. **A-C**. Mitotic progenitors in the cortical neurogenesis of mice and rats (Days 1–7) are immunolabeled with phospho-histone H3 antibody. Mitotic RG and IP are separately quantified in the ventricular zone (VZ, arrows in **A**; quantification in **B**) and in the subventricular zone (SVZ, parenthesis in **A**; quantification in **C**). **D**. Cells in the G2/S phase are labeled with EdU administered one hour before sacrifice. **E**. Total number of EdU-labeled cells during the neurogenic period is quantified. **F**. Double-labeled EdU and Tbr2-positive cells, identified as IP, are quantified across neurogenic stages. Statistical analysis: Time points with statistically significant differences (p < 0.05) are indicated by asterisks (**B**, **C**, **E**, **F**; Student’s t-test).

Despite the overall conservation, we noted a minor increase in SVZ mitosis in rats compared to mice (**Figure 5C**). Rats exhibited a relatively higher abundance of IPs in the G2/S phase, co-labeled by EdU and Tbr2, throughout the neurogenic period (**Figure 5F**). This suggests that indirect neurogenesis plays a more prominent role in rats than in mice, not limited to the early DL production phase but extending across the entire neurogenic period. In conclusion, differences in cell cycle kinetics and indirect neurogenesis are unlikely to be the primary factors driving the species-specific variation in DL neuron production. Instead, the temporal regulation of progenitor neurogenic competence, which determines the duration of the DL production phase, provides a more plausible explanation for the greater number of DL subtypes observed in rats (**Figures 3 and 4**).

### Non-uniform correspondence of temporal progression in cortical RG progenitors in mice and rats revealed by scRNAseq

To investigate the molecular basis of the species-specific temporal dynamics of cortical neurogenesis, especially the elongated period of DL production in rats, we performed comparative scRNAseq analysis of neurogenic progenitors. Using the Chromium X system (10x Genomics), we obtained high-quality transcriptome profiles for 14,290 cells from rat cortices at Days 1 (E14.5), 3 (E16.5), and 5 (E18.5) of neurogenesis (**Figure 6A**). After quality control, we integrated these profiles with previously published mouse cortical cell data from corresponding neurogenic stages(*36*) (**Figure S8**). Uniform Manifold Approximation and Projection (UMAP)(*37*, *38*) identified 16 distinct cell clusters annotated with commonly used cell type marker genes (**Figure 6B-C and Figure S8C-E**). All major cortical cell types were represented, with no cluster dominated by a single species or developmental stage, indicating clustering was driven by cell type differences rather than species-specific biases or technical artifacts (**Figure 6C and D**).

**Figure 6.**
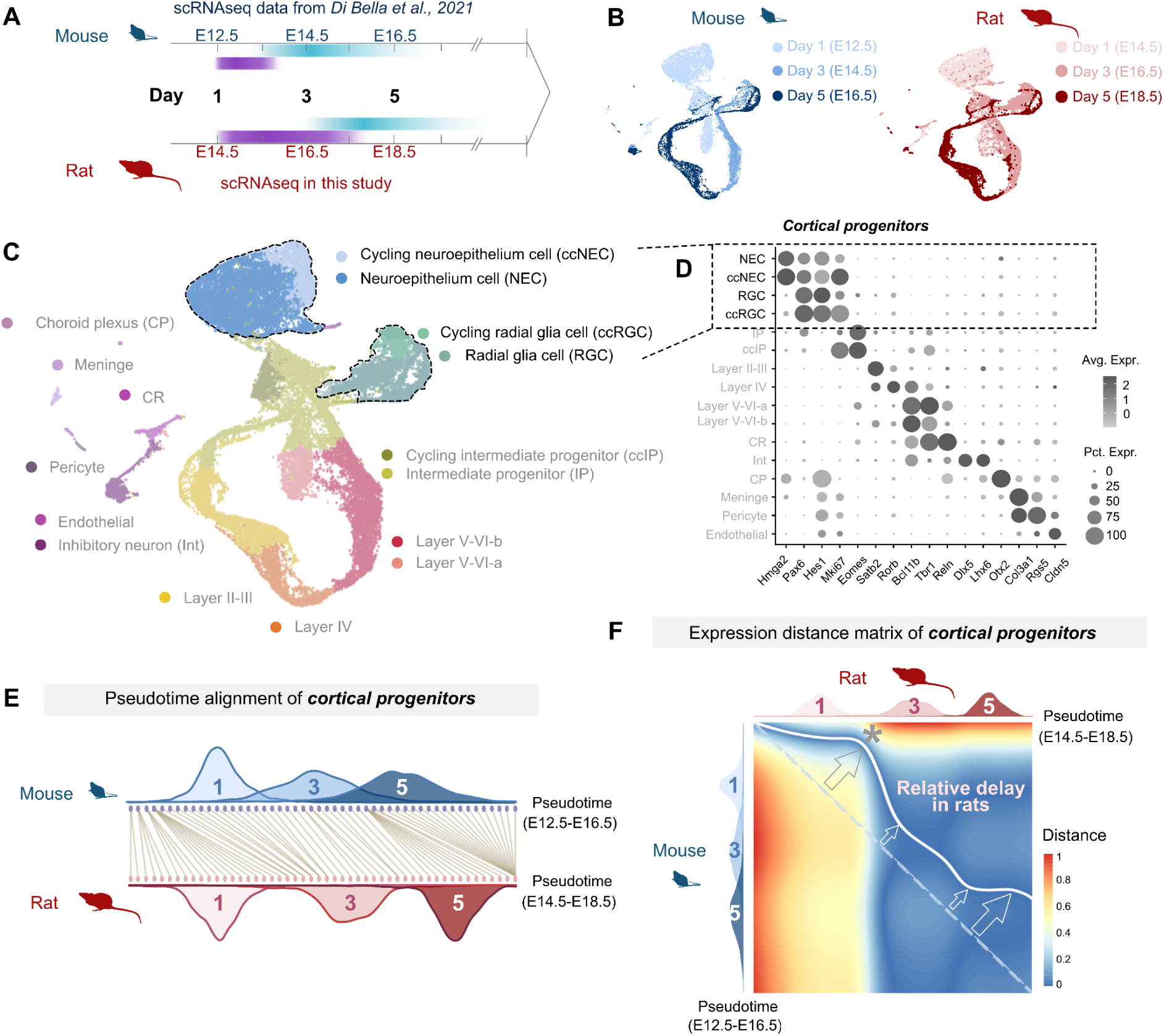
Temporal progression of progenitor aging in the mouse and rat cortical neurogenesis revealed by scRNAseq. **A**. scRNAseq analysis of rat cortical cells collected at three neurogenic stages (Days 1, 3, and 5) was performed and compared with previously published mouse datasets from corresponding stages. **B**. UMAP visualization of cortical cells from three developmental stages in mice and rats. **C**. UMAP representation of cortical cells from both species, illustrating the clustering of major cell types. **D**. Cell type-specific marker genes used for annotation, ensuring consistent identification across species. **E**. Alignment of mouse and rat progenitor cells based on their pseudotime scores. Trajectories in each species were divided into 50 blocks ("Pseudocells") for pairwise distance calculations using dynamic time warping. Histograms display the distribution of cells from sampling stages (Days 1, 3, and 5) along the trajectories. **F**. Distance matrix of mouse and rat progenitor cells based on "progenitor age gene" expression. Cells were aligned by pseudotime scores along species-specific trajectories. The optimal alignment is indicated by the solid white line, which deviates from the diagonal dotted line, reflecting delayed developmental progression in rats compared to mice. Inflection points (open arrows) highlight significant changes in relative developmental tempo, particularly in the early phase (asterisk).

We examined temporal gene expression changes in progenitors, as neuronal laminar fate is determined by the progenitor’s developmental age. Using a machine learning-based ordinal regression model(*39*), we identified stage-specific "progenitor age genes" (**Table S3** and **Figure S9A-F**), which showed significant overlap between species and with genes identified in previous studies of cortical neurogenesis(*40*, *41*). Gene ontology (GO) analysis revealed enrichment for terms such as "positive regulation of Wnt signaling pathway" (**Figure S9G**). To quantitatively compare temporal changes between species, we aligned progenitors by developmental age using a pseudotime analysis based on the expression profiles of progenitor age genes with cellAlign algorism(*42*) (**Figure 6E**).

Corresponding pairs of mouse and rat progenitors were identified by minimizing expression distance. This analysis revealed that most rat progenitors at Day 1 correspond to a smaller subset of mouse progenitors with the youngest pseudotime scores. This indicates that rat progenitors retained early-age expression profiles longer than mouse progenitors, which progressed through developmental aging more rapidly. In contrast, the later neurogenic phase progressed in parallel in both species (Days 1 and 3 in mice and Days 3 and 5 in rats), with exceptions at the end of neurogenesis. Non-linear correspondence in pseudotime changes was observed in more quantitative measurement, with the largest delay in rat progenitors occurring where Day 1 and Day 3 progenitors overlapped (asterisk in **Figure 6F**). In summary, the temporal progression of cortical progenitors exhibits species-specific non-uniform scaling, as revealed by single-cell resolution transcriptomics.

### Wnt ligand genes exhibit protracted expression in rat progenitor cells

To investigate the molecular mechanism of how the early neurogenic phase for DL neurons is extended in rats relative to mice, we performed an unbiased re-analysis of our scRNAseq dataset. We identified 431 highly variable genes (HVGs) with dynamic expression in RG and applied Weighted Gene Co-expression Network Analysis (WGCNA), which resolved four temporally defined modules (**Figure 7A, Table S3**). Modules 1 and 2 showed conserved trajectories across species: module 1 increased modestly with developmental progression, whereas module 2 declined monotonically. By contrast, modules 3 and 4 exhibited clear species divergence. Module 3 was a subset of module 4 with a related but distinct profile. In mice, module-4 genes decreased steadily over time; in rats, they remained high early and decreased later, mirroring the species-specific timing of DL neuron production. GO enrichment linked module 1 to microtubule binding, module 2 to pallial development, and module 4 to Wnt signaling pathway; module 3 showed no significant enrichment.

**Figure 7.**
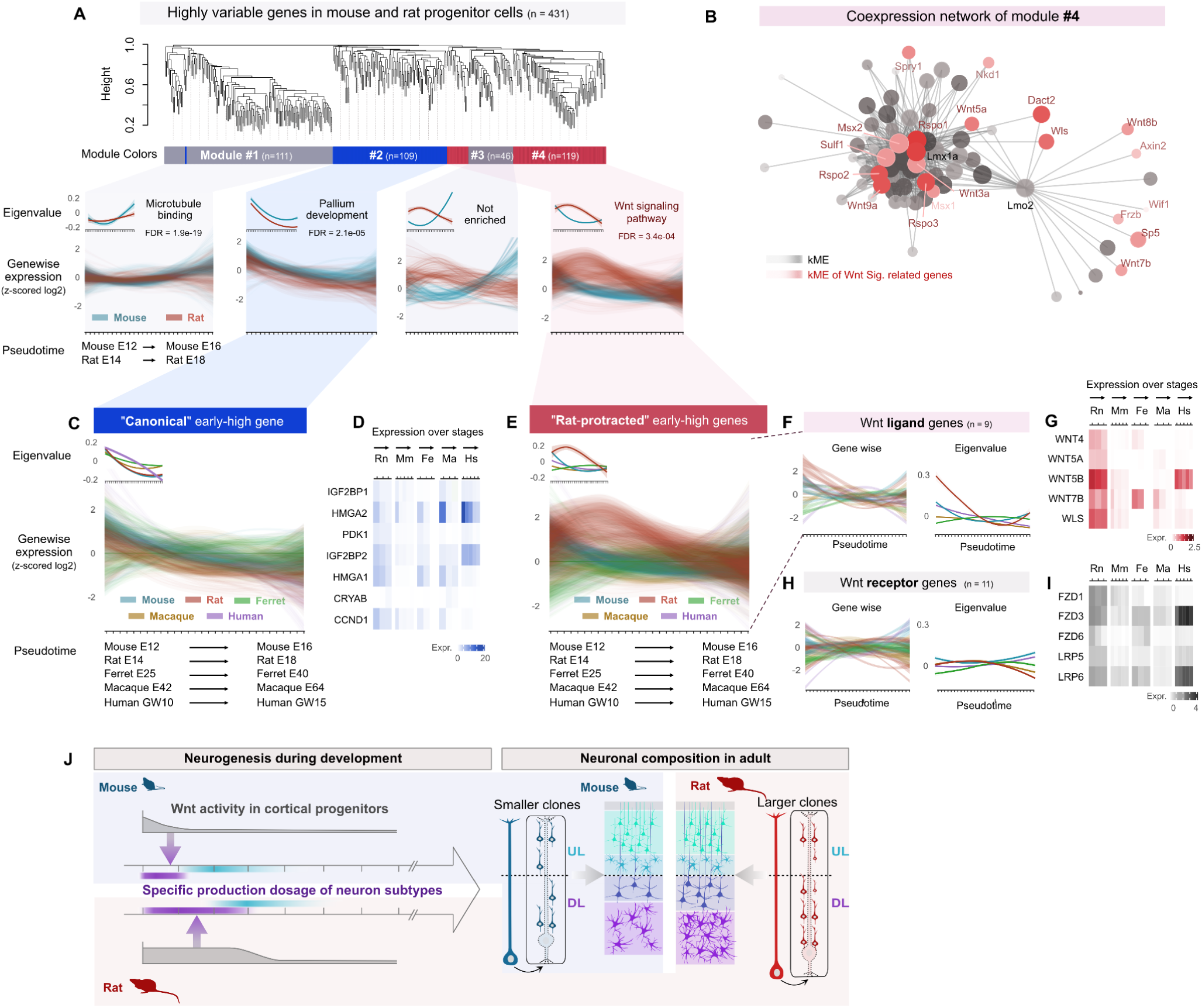
Protracted expression of Wnt ligand genes in rat cortical progenitors. **A.** Weighted gene co-expression network analysis (WGCNA) of highly variable genes (n = 431) in mouse and rat radial glia progenitors (RGCs) identifies four co-expression modules. Gene expression values were log-normalized and z-scored across pseudotime to show temporal dynamics (see Methods). Module eigenvalues (first principal component of each module’s gene expression matrix) and the top Gene Ontology (GO) term (lowest FDR-adjusted P-value) are indicated in the upper left and right of each plot, respectively. **B.** Co-expression network for module 4. The top 300 connections are shown, with edge shading representing module membership (kME) values. Genes associated with the Wnt signaling pathway are highlighted in pink to red based on kME. **C-D.** Expression dynamics of the "canonical" early-high module genes (n = 109) across developing somatosensory cortex progenitors in five mammalian species. **(C)** Log-normalized and z-scored expression along the pseudotime axis, highlighting temporal patterns. **(D)** Absolute expression values across actual developmental stages in each species. **E.** Expression dynamics of the "rat-protracted" early-high module genes (n = 111) across five mammalian species, as in (C and D). **F–I.** Temporal dynamics and absolute expression of Wnt ligand genes from module 4 (n = 9; **F** and **G**) and canonical Wnt receptor genes (n = 11; **H** and **I**) across progenitors in five mammalian species, as in (C and D). **J**. Schematic illustration summarizing this study.

Canonical Wnt signaling is a key regulator of the temporal program of corticogenesis and is elevated early, declining as development proceeds in both mouse and human(*39*, *41*). Experimentally increasing Wnt signaling in RG biases production toward DL subtypes, whereas reducing it shifts fate toward UL subtypes(*43–46*). Given the necessity and sufficiency of Wnt signaling for time-dependent subtype specification, we interrogated module 4 in detail. Its co-expression network contained transcription factors, multiple Wnt ligands, and pathway components such as *Axin2* and *Wls*, but notably lacked canonical Wnt receptor genes (**Figure 7B**). Fluorescent *in situ* hybridization using RNAscope confirmed significantly elevated *Axin2* expression in rat RG throughout neurogenesis compared with mouse, consistent with our scRNAseq results. Because *Axin2*, a standard readout of Wnt activity, displayed divergent dynamics between species (**Figure S10**), we hypothesized that species-specific Wnt kinetics underlie the different durations of DL neurogenesis.

To assess whether the mouse- or rat-like pattern is ancestral, we integrated publicly available single-cell datasets from human(*47*), macaque(*48*), and ferret(*49*) with our mouse and rat data. Across all five species, module-2 genes such as *Hmga2* and *Igf2bp2* (*IMP2*) decreased consistently with developmental progression (**Figure 7C–D**). In contrast, module-4 genes, particularly Wnt ligands, showed a rat-specific trajectory in RG: other mammals exhibited a rapid decline, whereas rats maintained high expression for a prolonged period (**Figure 7E–G**). Wnt receptor genes showed no clear temporal change in any species (**Figure 7H–I**). Elevated Wnt-ligand expression was specific to RG and was not observed in other cell types, including IP (**Figure S11**). *Lmx1a*, a master regulator of the cortical hem(*50*), a signaling center expressing multiple Wnt ligands, is highly expressed in RG progenitors, specifically in rats (**Figure S12**). This suggests a potential mechanism for the rat-specific induction of Wnt ligands in these cells. These findings indicate that sustained Wnt-ligand expression in RG is exceptional among the species examined and likely arose in the rat lineage.

Together, the data suggest that, after diverging from mice, rats evolved elevated and prolonged expression of Wnt-ligand genes specifically in RG. This likely maintained population-wide Wnt signaling through auto- and paracrine ligand supply, extending the DL neurogenic window and contributing to rat-specific expansion of deep cortical layers (**Figure 1A and Figure 7J**).

## Discussion

This study performed a comparative anatomical analysis of layer architecture within the primary somatosensory cortices of mammals. In contrast to the notable expansion of UL in primates, the rodents exhibit thicker DL (**Figure 1A**). Rats, in particular, exhibit an exceptionally high abundance of DL subtypes, even when compared to their close evolutionary relatives, mice.

Histological assessments revealed species-specific variations in cortical excitatory neuron composition: rats exhibit significantly more DL neurons per cortical column than mice, while UL neuron numbers remain comparable between the two species (**Figure 1**). Clonal analysis demonstrated that rat RG progenitors produce more DL neurons than those in mice, suggesting that different progenitor activity drives these species-specific differences (**Figure 2**). Birthdating assays further indicated an extended phase of DL neurogenesis in rats, followed by a conserved UL generation phase (**Figures 3 and 4**). Our scRNAseq analysis revealed that rats maintain the early progenitor state associated with DL neuron production for a longer period than mice. This was evidenced by the sustained expression of early-biased progenitor age genes (**Figure 6**). More detailed analysis revealed that rat RG progenitors exhibit significantly elevated and sustained expression of Wnt-related genes, most notably the genes encoding Wnt ligands, than the counterparts in mice. The integrated analysis with the data of other mammalian species demonstrated that the situation in rats is exceptional (**Figure 7**). These findings suggest that rats extend DL neurogenetic phase by prolonging Wnt high period, reflecting a species-specific, non-uniform scaling of an evolutionarily conserved neurogenic program (**Figure 7J**).

### Evolutionary adaptations in DL neuron composition

Our study demonstrates that mice and rats have undergone distinct evolutionary adaptations in the abundance of deep-layer (DL) neurons. This pattern contrasts with the human lineage, where cortical expansion is characterized primarily by an increase in upper-layer (UL) neurons (**Figure 1A**). In rats, all DL subtypes examined—including Tbr1-, Ctip2-, and Satb2-expressing neurons—were more abundant than in mice, whereas UL neuron numbers remained largely conserved (**Figure 1**).

This finding is consistent with anatomical observations: the rat corticospinal tract, derived from DL neurons, is considerably larger than its counterpart in mice, despite only minor differences in the thickness of the corpus callosum, which originates from UL neurons (**Figure S2**). An increase in Ctip2-expressing CSPNs in layer 5 may be linked to the larger body size of rats and their greater demands for motor control. Furthermore, although UL neuron abundance was comparable between species, rats exhibited a higher number of Satb2-positive neurons within the DL. This suggests an expansion of descending outputs from cortico-cortical circuits. Notably, Satb2-positive DL neurons have dual projections to both callosal and brainstem targets, in contrast to UL neurons, which project exclusively to callosal pathways(*51*, *52*). Finally, we note that the cortical surface is substantially more expanded in rats compared with mice. This expansion likely reflects a greater degree of progenitor amplification prior to the onset of neurogenesis. While the overall DL-to-UL ratio remains unchanged, this early amplification increases the total number of cortical neurons that contribute to axonal projections, thereby enhancing the output capacity of rat cortical circuits.

Further studies on species-specific DL neuron subtypes could clarify how anatomical differences influence functional and cognitive capabilities. The diversity of DL projection targets is also of interest. Our axon labeling experiments primarily classified projections as intracortical or extracortical (**Figure S7**). More detailed mapping of DL subtype projection patterns in rats(*51*), as has been done in mice(*53*, *54*), could elucidate interspecies differences in extra-cortical projectome. The potential behavioral consequences of varying DL neuron abundance and complexity remain to be explored. While comparative behavioral studies in mice and rats are important, a more precise understanding could be gained from genetically modified mice with specific alterations in DL abundance(*55*). This raises the broader question of whether cortical neuronal composition reflects evolutionary adaptations to distinct ecological niches. Rodent species occupy diverse habitats, and their brain architectures likely facilitate survival in these environments(*56*, *57*). Investigating how specific cortical features enhance evolutionary fitness would provide valuable insights.

### Temporal regulation of cortical neurogenesis by canonical Wnt signaling pathway

Canonical Wnt signaling is a key temporal regulator of cortical neurogenesis(*43–45*). In both mice and humans, pathway activity is high during early neurogenesis and diminishes as development proceeds(*39*, *41*). Experimental perturbations in mouse RG progenitors demonstrate causality: elevating Wnt signaling favors production of DL neurons, whereas reducing it promotes UL identities(*43*, *44*, *46*). Heterochronic transplantation further supports this temporal control(*45*). Late-stage RG placed into an early Wnt-rich VZ environment reacquire DL potential, and this “rejuvenation” requires intact Wnt signaling.

Our data, together with prior work(*45*), indicate that Wnt signaling level in RG is governed primarily by the developmental dynamics of Wnt ligands rather than by their receptors (**Figure 7**). Across stages, Wnt receptor expression is comparatively stable, preserving the ability of RG to sense Wnt inputs, whereas ligand expression changes markedly. Consequently, when late-stage RGs encounter an early environment with abundant Wnt ligands, their constant receptor repertoire permits robust pathway activation and restoration of early-born DL output(*45*). We propose a model where the concentration of Wnt ligands within the ventricular progenitor niche establishes the temporal window for RG neurogenic competence. Our proposed model suggests that the concentration of Wnt ligands within the niche, initially high and subsequently decreasing, determines the timeframe during which RG cells maintain their neurogenic competence. Building on this framework, our comparative analysis suggests that species differences in the time course of Wnt ligand availability in RG can help explain interspecies variation in neuronal subtype balance (**Figure 1A** and **Figure 7**).

Among the five mammals examined, the rat is a clear outlier: RGs maintain unusually prolonged, high expression of Wnt ligand genes (**Figure 7**). By analogy to experimental perturbations in mice(*44*), this sustained ligand milieu likely contributes to the extended production of DL neurons in rats. What sustains high Wnt levels specifically in rats? Known sources of cortical Wnt ligands include the cortical hem and postmitotic neurons(*58–62*). The hem, the dorsomedial telencephalic organizer, expresses multiple Wnt ligands and is under control of the master regulator *Lmx1a*: inhibiting *Lmx1a* suppresses the expression of several Wnt ligands, while forced *Lmx1a* expression in the future hippocampal region can induce *Wnt3a*(*50*). In the midbrain, *Lmx1a* and *Wnt1* engage in a positive regulatory loop(*63*). Consistent with a strong medial Wnt output, the pathway readout *Axin2* is highest medially and tapers laterally(*64*, *65*). In parallel, early-born subplate and DL neurons express ligands such as *Wnt7b*, which can bias newly generated neurons toward DL fates(*61*, *62*).

Extending these observations, our analysis shows that rat RG themselves express multiple Wnt ligands together with the essential secretion factor *Wls* (**Figure 7**). In rats, a co-expression module that remains elevated comprises Wnt ligands, *Wls*, and *Lmx1a*, recapitulating a hem-like regulatory architecture and implicating *Lmx1a* as a potential upstream driver of Wnt ligand expression within RG. Notably, only in rats did a substantial subset of cortical RG co-express *Lmx1a* and *Lhx2* (**Figure S12**), even though in mice the cortical fate selector *Lhx2* suppresses *Lmx1a*, a master regulator of the cortical hem(*66*). In our comparative transcriptome, the other mammals examined showed virtually no RGs with this co-expression, suggesting a rat-specific co-option of an *Lmx1a*–Wnt ligand cascade in cortical RG, potentially via relief of *Lhx2*-mediated repression of *Lmx1a*. Dissecting the regulatory logic that induces *Lmx1a* in rat cortical RG, and testing the requirement of *Lmx1a* for extending the DL production window, will be informative. Together, these findings support a model in which a sustained, RG-intrinsic and niche-derived Wnt ligand environment, potentially orchestrated by *Lmx1a*, prolongs DL neurogenesis in rat and helps explain species-specific differences in cortical neuron subtype composition.

### Evolvability of the mammalian cortical neurogenesis

Our study highlights a distinction between conserved and evolved components of the mammalian neurogenetic program. The early-to-late pattern of cortical neurogenesis, regulated chiefly by canonical Wnt signaling, is highly conserved across mammals(*11*, *13*, *14*, *67*, *68*) and beyond(*69–71*), suggesting that this developmental process has been maintained due to robust constraints(*72*, *73*). Despite its apparent rigidity, our comparative analysis of closely related species revealed that subtle adjustments of heterochronic regulation can drive species-specific cortical neuron compositions without fundamentally altering the overall developmental framework. Specifically, the genetic system activating Wnt ligand expression, presumably driven by *Lmx1a*, was co-opted in early RG progenitors during the evolutionary specification of rats. Consequently, the temporal dynamics of Wnt signaling activity was fine-tuned to prolong DL subtype-producing period. This non-uniform scaling of temporal progression mirrors differences observed in other developmental processes, such as cell differentiation trajectories during gastrulation in mice and rabbits(*74*) and brain development in eutherians and marsupials(*75*). However, it contrasts with the relatively uniform elongation of developmental processes observed in humans compared to mice, as evidenced by studies on cortical neuron maturation(*76–78*), spinal motoneuron generation(*79*), and somitogenesis(*80–82*). Future comparative studies across a broader range of species could help clarify the balance between highly constrained components of the neurogenetic program and its more adaptable elements. Such insights would illuminate how evolutionary diversification optimized cortical architectures for species-specific functional and ecological needs.

## Limitations of the study

This study focuses on the generation of cortical excitatory neuron subtypes in mice and rats, leaving several critical areas unexplored. For instance, we did not examine progenitor amplification preceding neurogenesis in detail, despite notable differences in cortical neuroepithelial sheet size at the onset of neurogenesis. Investigating how neuroepithelial expansion varies between species is crucial for understanding the determinants of species-specific cortical volume and surface area.

Additionally, our study did not address other cell types, such as inhibitory neurons and glia, in depth. For example, rats exhibit a higher number of proliferating Olig2-positive glial progenitors during early postnatal periods compared to mice (**Figure S4C**). These differences in postnatal cell division events may align with the large deviations in developmental tempo observed in progenitors during the late phase of neurogenesis, as revealed by scRNA-seq (**Figure 6F**). Such findings suggest that rats may possess an increased abundance of glial progenitors, potentially resulting in greater diversity and quantity of glial cells.

## Supporting information

Key Resource Table

## Acknowledgements

We are grateful to our colleagues in the lab and common laboratory facilities in the WINGS-LST graduate program at the University of Tokyo. We thank Mr. Naoto Ohte, Drs. Ryohei Iwata, Noruhiko Yamamoto, Tatsumi Hirata, Tadashi Nomura, Kaoru Sugimura, Carina Hanashima, Akira Uematsu, Kenji Shimamura and Chiaki Ohtaka-Maruyama for insightful suggestions for the project and the manuscript. Computations were partially performed on the NIG supercomputer at ROIS National Institute of Genetics. The large Japanese field mouse are kindly provided by Kobe animal kingdom.

## Author contributions

I. K. S. co-ordinated the project and helped to design, perform and interpret experiments. Y. Y. Y. and I. K. S. designed and performed the experiments and interpreted the results with the help of X. D. S., P. R. and T. R.. Y. Y. Y., X. D. S., D. S., T. N., M. B., and Y. K. performed the experiment of single-cell transcriptome. X. D. Z. led the analysis of scRNAseq data. K. E. contributed to supervision. Y. Y. Y., X. D. S., and I. K. S. analyzed the data and prepared figures. Y. Y. Y., X. D. S., and I. K. S. wrote the paper. All authors read and approved the manuscript.

## Data, Code and Materials availability

All animals and unique/stable reagents generated in this study are available from the lead contact with a completed Materials Transfer Agreement.All data and the codes for analyzing the data in this study are available from the lead contact. Single cell RNA sequencing data of rat cortical cells are available in NCBI Gene Expression Omnibus (GEO: GSE287210).

## Supplementary Materials

### Materials and methods

#### Animals

All animal procedures were approved by the Institutional Safety Committee on Recombinant DNA Experiments and the Animal Research Committee of the University of Tokyo. ICR mice and WistarST rats were purchased from SLC Japan. Mice and rats were housed in cages with bedding (Avidity Science, TEK-FRESH) and provided constant access to food (Nippon Crea, Rodent Diet CE-2) and water. The animal facility was maintained at a temperature of 23 ± 2°C, a humidity of 50 ± 10%, and a 12-hour light/dark cycle. Plug day was defined as embryonic day (E)0.5, and the day of birth as postnatal day (P)0. Data from all embryos were pooled without discrimination of sex, given the difficulty of determining sex identity at embryonic stages. Rabbit and the large Japanese field mouse were obtained in the previous project in Tokyo Metropolitan Institute of Medical Science(*83*). Ferrets were provided by SLC Japan and Marshall Farms (North Rose, NY), and Hartley guinea pigs were obtained from SLC Japan in the previous project in Kumamoto University(*84*).

#### Nissl staining

We performed Nissl staining following the protocol provided with the kit (Thermofisher, # N21483). Briefly, cryosections were rehydrated for ≥40 min in 0.1 M PBS (pH 7.2), permeabilized for 10 min in PBS with 0.1% Triton X-100, and rinsed. NeuroTrace was diluted in PBS (100×), applied to cover the tissue (800 µL/slide) for 20 min, removed, and sections were washed in PBS/0.1% Triton X-100 (10 min), PBS (2 × 5 min), then PBS for 2 h at room temperature or overnight at 4 °C. Sections were mounted with DAKO glycerol mounting medium (Cat# C0563) and stored in the dark at 4 °C. Human and macaque (primary somatosensory cortex, face- and neck-related subregions) as well as adult mouse (somatosensory barrel field) brain sections were obtained from the Allen Brain Reference Atlas, with detailed image identifiers listed in the corresponding supplementary figures. Mouse, rat, ferret, the large Japanese field mouse, guinea pig, and rabbit brain sections (adult somatosensory barrel field) were prepared in-house.

#### Preparation of retroviruses

HEK293T cells were seeded to reach 80–90% confluency on a 10 cm dish. After 24 hours, the cells were transfected with a mixture containing: 0.85 µg pCMV-VSV-G (Addgene #8454), 9.90 µg pUMVC (Addgene #8449), 8.25 µg pRC-2A-EGFP (kind gift from Dr. Pierre Vanderhaeghen and Ryohei Iwata) or pRC-mCherry (prepared in this study). The transfection was performed in 1200 µL of Opti-MEM supplemented with 60 µL of X-tremeGENE. After 24 hours, the medium was replaced, and the cells were cultured for another 24 hours. The supernatant was then filtered through a 0.45 µm filter, and retroviral particles were concentrated by centrifugation at 25,000 rpm (82,700 g) at 4°C for 2 hours using a 20% sucrose/PBS cushion. The viral pellet was resuspended in 200 µL of cold DPBS, aliquoted in 20 µL, and stored at -80°C. Viral titers were checked using HEK293T cells for each batch.

#### Retroviral injection to animals *in utero* for clonal tracing

Pregnant females were anesthetized with isoflurane (Viatris, Cat# 114133403), and the uterine horns were exposed under sterile conditions. One microliter of retroviral solution mixed with 1% Fast Green was injected into the fetal lateral ventricles using a heat-pulled capillary. After injection, the uterine horns were returned to the abdominal cavity, and the incisions were sutured. Mice were kept on a heating plate until fully recovered.

### *in utero* electroporation

*in utero* electroporation was performed with modifications from established protocols(*85*, *86*). Timed-pregnant rats (E16.5) or mice were anesthetized with isoflurane. Plasmid solutions (1–1.5 mg/mL DNA) were injected into the embryonic lateral ventricles using heat-pulled capillaries.

Electroporation was conducted using tweezer electrodes (Nepa Gene, Cat# CUY650P5) connected to a NEPA21 Type II electroporator (Nepa Gene) with the following settings: Rats: Voltage 50 V, pulse duration 50 ms, interval 950 ms, 4 pulses. Mice: Voltage 50 V, pulse duration 50 ms, interval 950 ms, 6 pulses. Embryos were returned to the abdominal cavity, and the mothers were sutured and placed on a heating plate until recovery. Brains were collected from electroporated pups at P7.

### DNA constructs

Membrane-anchored Achilles was amplified by PCR using primers with a membrane targeting signal sequence from the template plasmid (Addgene #153528) and subcloned into a pCAG plasmid.

### Histology and immunostaining

Immunostaining was performed as described previously(*69*, *86*). Mouse embryos were collected after electroporation and perfused transcardially with ice-cold 4% paraformaldehyde (PFA) in PBS. Dissected brains were soaked in 4% PFA overnight at 4°C and sectioned into 100 µm slices using a vibratome (Leica, Cat# VT1000S). Slices were washed with PBST (PBS with 0.1% Triton X-100) three times and incubated in blocking solution (PBS with 0.3% Triton X-100 and 3% donkey serum) for 30 minutes. Brain slices were then incubated overnight at 4°C with primary antibodies (**Table S1**). After three PBST washes, slices were incubated with secondary antibodies (**Table S1**) for 2 hours at room temperature. After additional PBST washes, slices were mounted on slide glasses using DAKO glycerol mounting medium (Cat# C0563). Imaging was performed with a confocal microscope (Olympus FV3000), and images were processed using Fiji/ImageJ software(*87*).

### Clonal analysis

For clonal analysis, cortical sections were immunostained with anti-GFP and anti-RFP antibodies and imaged sequentially from rostral to caudal using a Slide Scanning Microscope (Keyence, Cat# BZ-X800). Clones were identified as sparse, spatially isolated cell clusters, typically separated by at least 500 µm. Cortical layers (UL vs. DL) were delineated using DAPI staining to define laminar boundaries, and the positions of labeled cells were recorded. Clones were categorized into UL-only, DL-only, or Mixed (UL and DL) types based on the laminar positions of their constituent cells. Clonal lineages with fewer than three cells or more than 30 cells were excluded from quantitative analyses.

### Neuronal birthdating

Birthdating analysis was performed as described previously(*88*). Pregnant animals were injected intraperitoneally with a thymidine analog, 5-ethynyl-2’-deoxyuridine (EdU, Fujifilm; Cat# 052-08843), at 45 mg/kg body weight at designated time points. Brains were collected at P7, and 100 µm coronal sections were prepared using a Vibratome (Leica, Cat# VT1200S). Sections were immunostained with anti-NeuN and anti-Olig2 antibodies to visualize neurons and glial progenitors. EdU incorporation was detected following the Invitrogen EdU detection kit protocol (Click-iT EdU Cell Proliferation Kit, Cat# C10637). EdU-positive cells were counted in cortical sections of 3–4 pups imaged with a confocal microscope (Olympus FV3000).

### Tissue clearing and axonal tracing

Tissue clearing was performed using the CUBIC trial kit (Tokyo Chemical Industry Co., Cat# C3942). Fixed samples were washed in PBS for 2 hours (repeated three times) and serially incubated in: 50% CUBIC-L solution for 12 hours at room temperature (RT)100% CUBIC-L solution for 72 hours at 37°C on a shaker. Samples were washed again in PBS for 2 hours (repeated three times), pre-treated in 50% CUBIC-R+ solution for 24 hours at RT on a shaker, and incubated in 100% CUBIC-R+ solution for 24 hours at RT on a shaker. Cleared brains were preserved in CUBIC-R+, and imaging was conducted with a confocal microscope (Olympus FV3000) in a 3.5 mm dish. Images were processed using Fiji/ImageJ software(*87*).

### Single cell transcriptomics

#### Sample preparation and sequencing

Rat embryonic brains were collected at E14.5, E16.5, and E18.5 and dissociated in ice-cold HBSS (Cat# H6648-500ML, Sigma) containing 0.1% BSA (Cat# 15260-037, Gibco). Tissues were digested in a 25 U/mL papain solution (Cat# 76216-50MG, Sigma) with 55 U/mL DNase (Cat# D4513, Sigma) and 0.5 mM EDTA for 20 minutes. After dissociation and concentration, cells were resuspended in HBSS with 0.04% BSA and processed immediately for single-cell GEM formation, cDNA amplification, and library construction using the Chromium Next GEM Single Cell 3’ Kit v3.1 (10x Genomics, PN-1000269) according to the manufacturer’s protocol. Libraries were quantified using an Agilent Bioanalyzer and sequenced on the NovaSeq6000 platform (Illumina) with an average read depth of 45,238 (E14.5), 42,669 (E16.5), and 44,504 (E18.5) reads per cell. Sequenced reads were aligned to the rat genome (mRatBN7.2) and processed with CellRanger v6.0.1. Noise reduction was performed using RECODE software(*89*).

#### scRNAseq analysis

Downstream analysis was conducted using Seurat v5(*90*). For quality control, cells were filtered using thresholds of nUMI>15000, nGene>15000, log10GenesPerUMI>0.9 and mitoRatio<0.25 to retain high-quality cells. To integrate rat scRNA-seq data with previously published mouse datasets(*36*), we normalized and extracted the 3,000 most variable genes using SCTransform and aligned cells between species with canonical correlation analysis(*91*). PCA dimensionality reduction was performed on integrated data, retaining the first 50 PCs. Cell clustering was conducted using the Louvain algorithm(*92*) with a resolution of 1.4. Cluster identities were assigned based on the top 50 differentially expressed genes (ranked by -log10 adjusted p-value, Wilcoxon Rank Sum test with Bonferroni correction), ensuring >30% cluster expression and a >0.5 log-fold change threshold, alongside canonical markers (**Figure 6C**).

#### Pseudotime age prediction and progenitor age genes

Pseudotime analysis followed established methods(*39*, *93*). L1-regularized ordinal regression was used to predict progenitor cell ages (neuroepithelial and radial glial cells). Analysis was restricted to the 3,000 integration genes to minimize batch effect and sequencing depth variations in the integrated datasets of rat and mouse cells(*36*). Highly variable genes were identified using the *scran* R package(*94*). We performed the *L1* penalized ordinal regression following the manual from the *psupertime* R package(*93*). After the cross-validation (*n_folds* = 10), the weight of the optimized model was used to rank the genes according to their ability to predict each progenitor cell age. Genes with non-zero coefficients in both mouse and rat models were classified as “progenitor age genes,” reflecting developmental temporal processes (**Table S3**).

#### Dynamic time warping-based alignment of pseudotime trajectories in mouse and rat

Dynamic time warping (DTW) of pseudotime trajectories was performed using CellAlign(*42*). Pseudotemporal trajectories for each species were equally spaced into 500 points. Pairwise Euclidean distances between ordered points were calculated using the shared progenitor temporal genes. A fixed-start/fixed-end constraint was applied, identifying the optimal alignment path to minimize overall distance. Gaussian weighting (winSz = 0.1) was applied to reduce noise.

#### WGCNA Analysis and Module Identification

Weighted gene co-expression network analysis (WGCNA)(*95*) was performed to identify co-expression modules associated with neurodevelopmental dynamics in radial glia progenitors.

Highly variable genes (HVGs) were first identified within the ventricular radial glia (vRG) cell subset from mouse and rat scRNA-seq datasets using the FindVariableFeatures function (selection.method = "vst", nfeatures = 2000) in Seurat (v5). The intersection of mouse and rat HVGs yielded 431 genes exhibiting consistent temporal dynamics in rodent progenitor cells. To focus on temporal patterns while mitigating inter-species magnitude differences, gene expression values were log-normalized from RNA assay and then z-scored across pseudotime trajectories (with start and end points aligned between species based on established developmental correspondence). WGCNA was implemented using the WGCNA R package (v1.73) with the following parameters: soft-thresholding power = 6 (selected based on scale-free topology fit), TOMType = "signed Nowick", and minModuleSize = 30 via the blockwiseModules function. This analysis identified four modules based on co-expression patterns (designated as modules 1–4, **Table S3**). Module eigengenes (defined as the first principal component of each module’s gene expression matrix) were visualized in the upper left of corresponding plots. Gene Ontology (GO) enrichment analysis was conducted for genes in each module using clusterProfiler (v4.6.2), with the most significant GO term (lowest FDR-adjusted P-value) displayed in the plot. Co-expression networks were visualized by selecting the top 300 connections ranked by module membership (kME) values.

#### Gene Expression Dynamics Across Five Mammalian Species

Building on scRNA-seq datasets from developing somatosensory cortex in rat and mouse(*36*), we incorporated three additional datasets from human(*47*), rhesus macaque(*48*), and ferret(*49*) to examine progenitor gene expression dynamics across species. Developmental stages were curated to span the equivalent period of mouse E12–E16 (rat E14–E18) using an established translational timing model(*96*), corresponding to gestational week (GW) 10–15 in human, embryonic day (E) 42–64 in macaque, and E25–E40 in ferret. Following subsetting to progenitor populations, pseudotime trajectories were inferred using an ordinal logistic regression model(*93*). Gene expression was log-normalized from RNA assay and z-scored across the pseudotime axis to capture temporal dynamics. To assess absolute expression levels while accounting for technical variability, values were retrieved from Seurat’s SCT assay and regressed against covariates ("nCount_RNA", "nFeature_RNA", "percent.mt", "CC.Difference") using the glmGamPoi method.

### Fluorescent *in situ* hybridization with RNAscope

Rat cortical tissues were cryosectioned at 20 µm thickness using Cryostat (Cat# CM3050S, Leica) and mounted on slide glasses (Cat# MAS-05, Matsunami Glass Ind.,Ltd) following the RNAscope protocol (UM323110, ACDBio). Sections were stored at -80°C until use. Preprocessing, probe hybridization, signal detection, and quantification were performed using the RNAscope Multiplex Fluorescent Detection Kit V2 (Cat# 32310, ACDBio).

### Quantification and statistical analysis

Statistical analyses were performed using GraphPad Prism 10 (GraphPad Software, Inc., USA) and Microsoft Excel (Microsoft Inc., USA). Experiments were repeated at least three times, and results are expressed as mean ± SEM. Student’s t-test: **Figures 1C, E; 5B-C,E-F**. Welch’s t-test: **Figures 2E, F; 3B; 4C-H**. Statistical significance thresholds and all quantitative results are summarized in **Table S2**.

**Figure S1.**
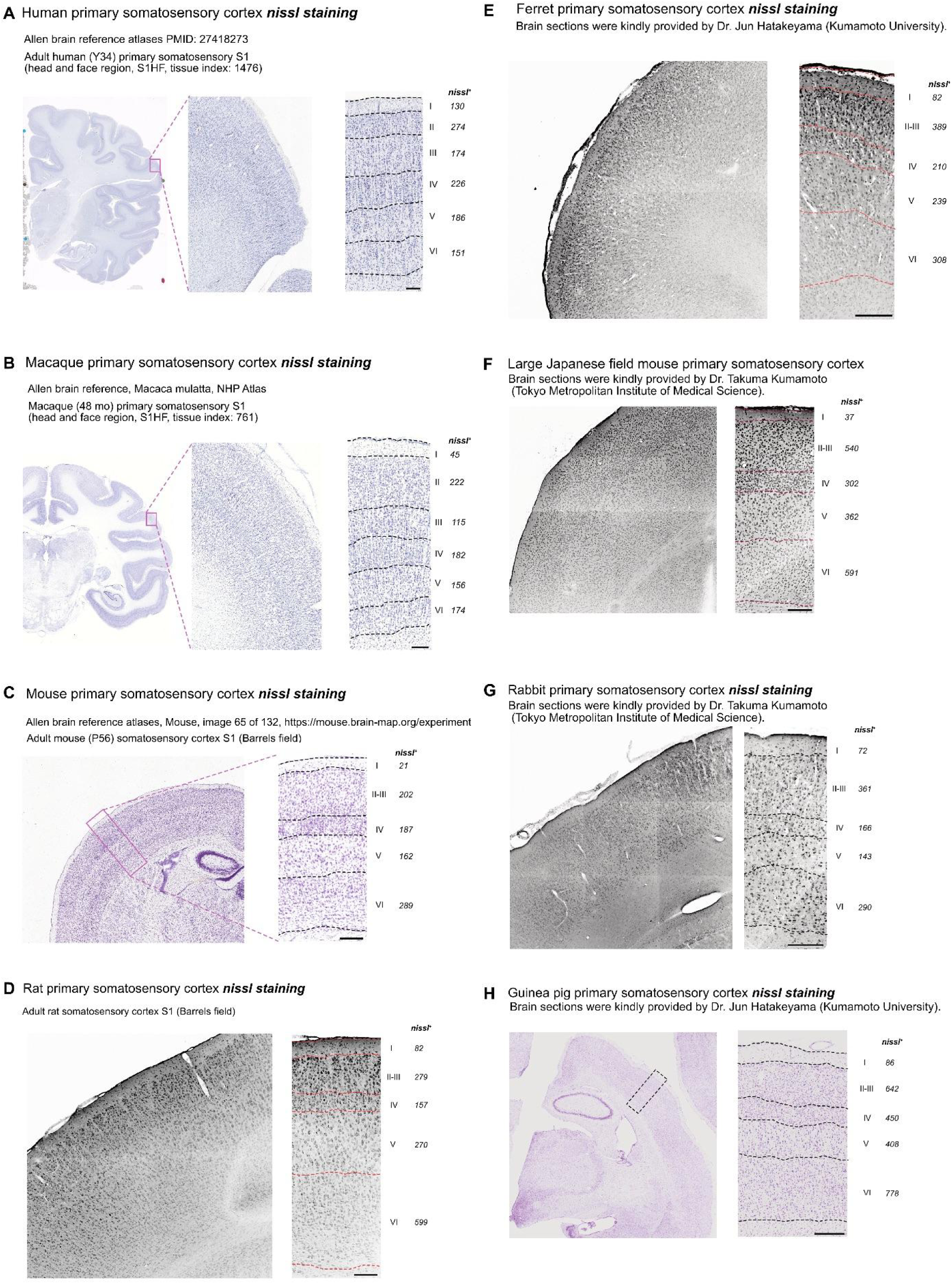
Comparative anatomy of somatosensory cortices of mammalian species. **A-H.** Coronal brain sections from adult animals (human, macaque, mouse, rat, ferret, large Japanese field mouse, rabbit, and guinea pig) illustrating the somatosensory cortex are presented in panels **A to H**. In rodents (mouse, rat, large Japanese field mouse, and guinea pig), sections were selected to highlight the barrel fields within the primary somatosensory cortex. For non-rodent species (human, macaque, ferret, and rabbit), where barrel fields are less distinct, regions corresponding to head and neck sensory input in the primary somatosensory cortex were curated as the closest homologous areas. Nissl staining was performed on brain sections from rat, ferret, large Japanese field mouse, rabbit, and guinea pig (see Materials and Methods for details). Human, macaque, and mouse brain sections were sourced from the Allen Brain Institute reference atlas (https://mouse.brain-map.org/static/atlas) to ensure anatomical consistency in the selected regions. Scale bars: 200μm **(A–H)**

**Figure S2.**
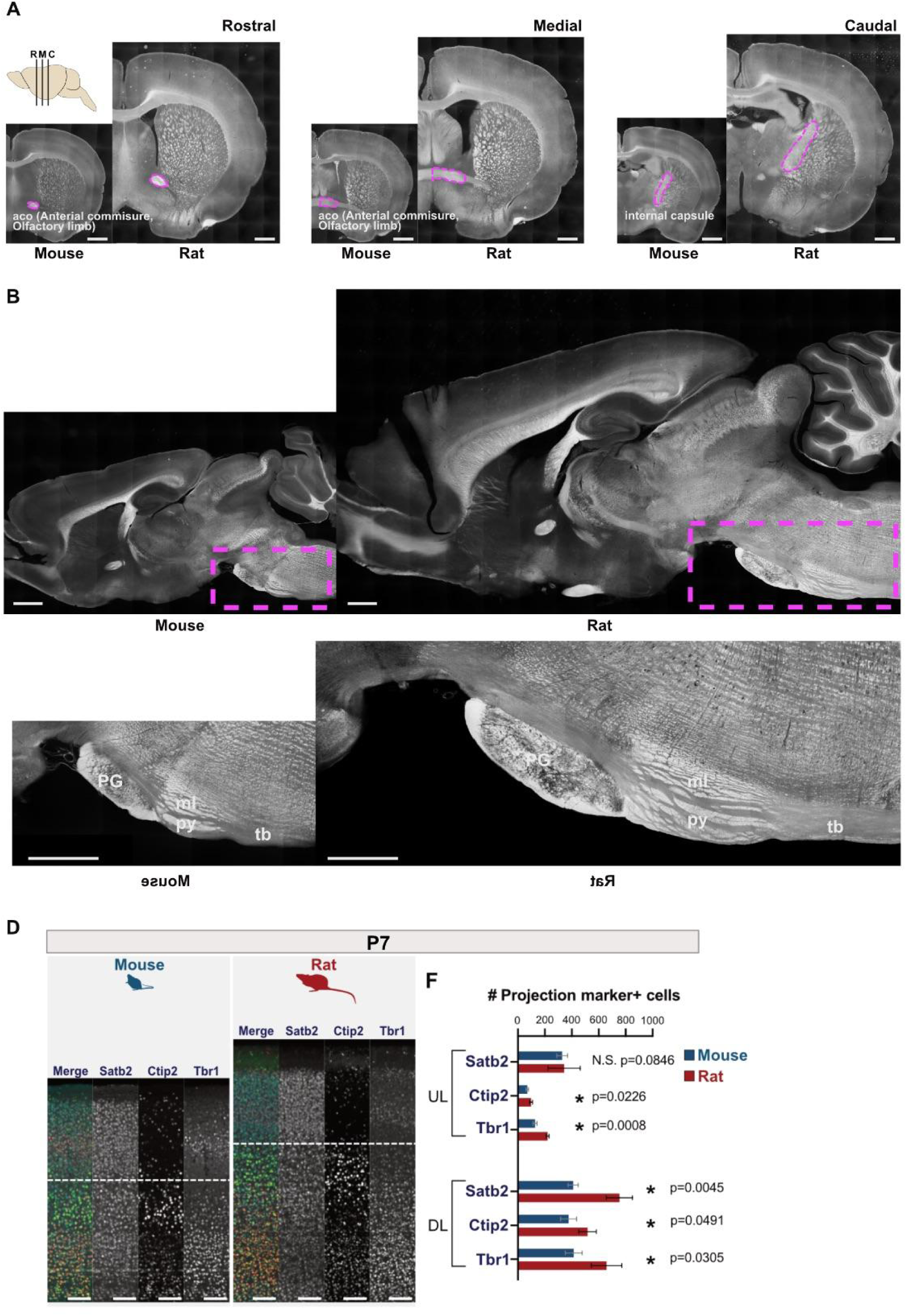
Intracortical and corticofugal projections as well as projection neuronal markers in mice and rats. **A.** Coronal sections of adult mouse and rat brains at rostral, middle, and caudal levels (R, M, C). Regions outlined with magenta dashed lines indicate the anterior commissure (ACO) and internal capsule (IC) in mice and rats, respectively. Axon-enriched areas appear whitish under transmitted light differential interference contrast (TD) imaging, clearly delineating the corpus callosum, ACO, and IC projections. **B.** Sagittal sections of adult mouse and rat brains visualizing corticospinal projections under TD imaging. Magenta dashed lines in panel **B** highlight corticospinal projections, with panel **C** providing an enlarged view of the corticospinal tract, including the pontine gray/nuclei (PG), medial lemniscus (ML), pyramidal tract (PY), and trapezoid body (TB). **D.** Immunohistochemical staining for projection neuronal markers (*Satb2, Ctip2,* and *Tbr1*) in the primary somatosensory cortex (S1) of mice and rats at postnatal day 7 (P7). White dotted lines demarcate the boundary between upper layers (UL) and deeper layers (DL). **E.** Quantification of Satb2-, Ctip2-, and Tbr1-expressing neurons in the UL and DL of mouse and rat S1 at P7. Data are presented as mean ± SEM (n = 3 samples per group); statistical significance was determined using Student’s t-test. Scale bars: 1mm **(A–C),** 100μm **(D).**

**Figure S3.**
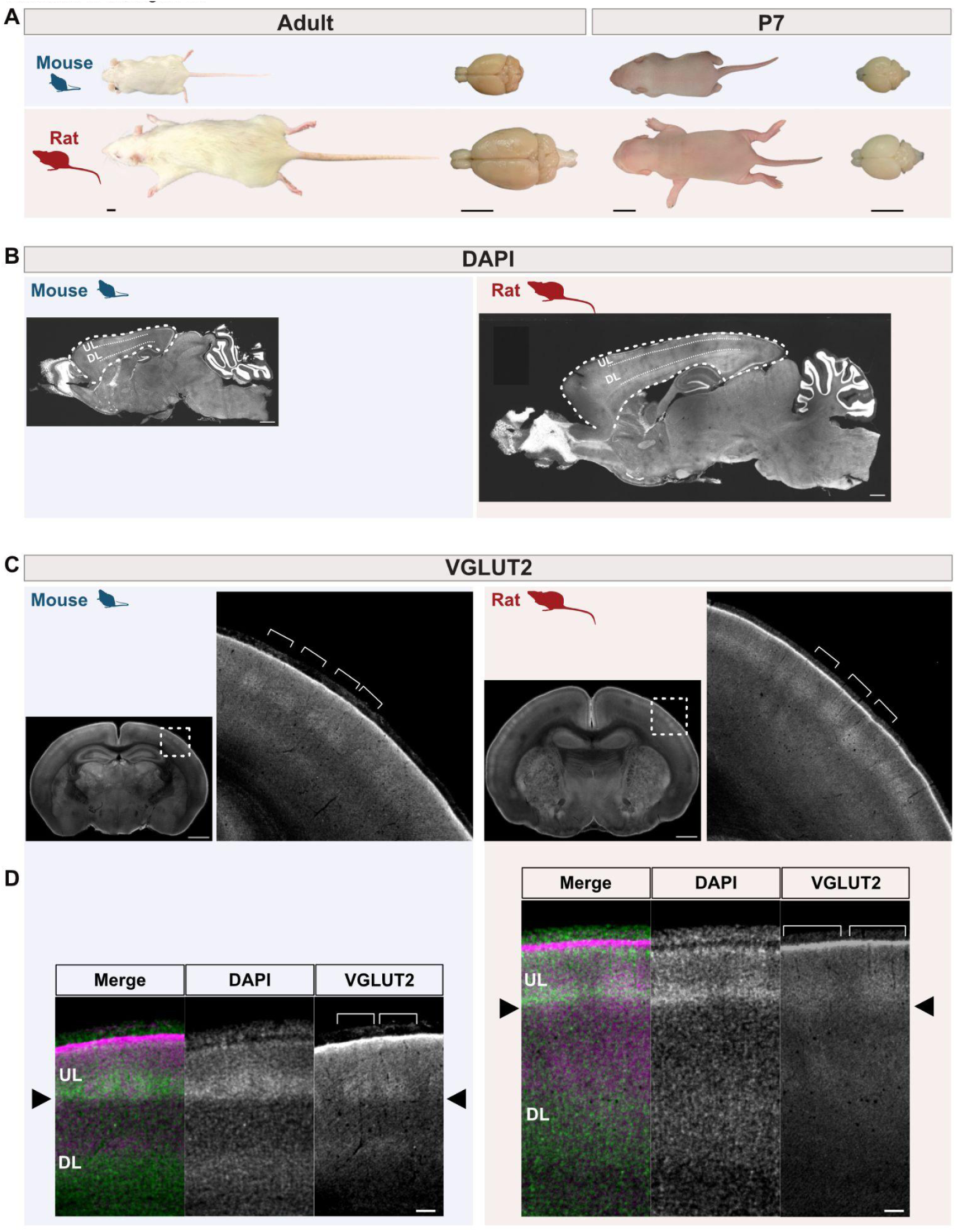
Morphological and histological comparison of mouse and rat cerebral cortices, related to Figure 1. **A**. Whole-body and brain size differences between mice and rats. **B**. Sagittal sections of mouse and rat brains at identical magnification. **C**. Analyses were conducted in the primary somatosensory field (S1) of both species, identified by the characteristic barrel structure of thalamic axon terminals visualized by VGLUT2 immunostaining. Magnified images highlight the structure within dotted rectangles. **D**. The border between UL (layer 2-4) and DL (layer 5 and 6) is defined by differences in cell density (DAPI staining) and the lower limit of VGLUT2-positive thalamic fibers (arrowheads). Scale bars: 1 cm (**A**), 1 mm (**B**), 1 mm (**C**), 100 μm (**D**).

**Figure S4.**
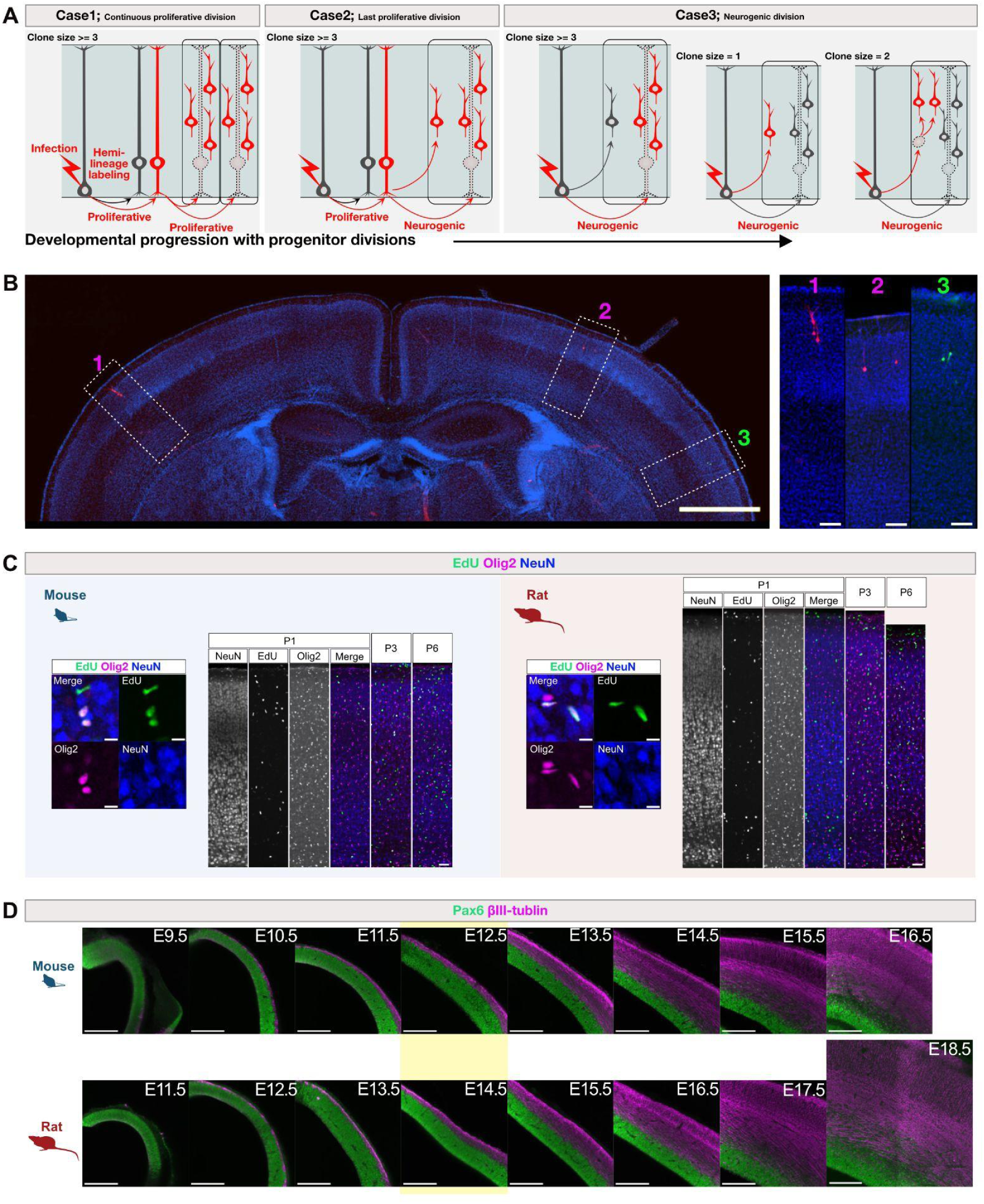
Clonal analysis of cortical RG progenitors in mice and rats from the onset to the end of excitatory neurogenesis, related to Figure 2. **A**. Schematic representation of hemi-lineage labeling via retroviral gene transfer, illustrating three scenarios of progenitor labeling. Cortical progenitors initially divide symmetrically to expand their population before transitioning to neurogenesis. Radial glial (RG) progenitors begin producing neurons by switching from symmetric proliferative divisions to asymmetric neurogenic divisions. Eventually, RG progenitors terminate producing neurons to begin gliogenesis. A clonal "unit" of neurons is defined as sister excitatory neurons that share a common progenitor origin and are produced during the first and subsequent neurogenic divisions of the mother progenitor (rounded boxes). **Case 1**: The retroviral vector infects a progenitor undergoing symmetric proliferative divisions. This results in multiple clonal units being labeled within the same cell cluster. **Case 2**: The retroviral vector infects an RG progenitor at its last proliferative division before initiating neurogenesis. In this ideal case, the entire clonal unit of sister neurons is labeled. **Case 3**: The retroviral vector infects an RG progenitor that has already started neurogenic divisions. In this scenario, some members of the clonal unit are missing from the labeled cohort, resulting in an incomplete clone. Alternatively, only one neuron (direct neurogenesis) or two neurons (indirect neurogenesis via intermediate progenitors) may be labeled.Given the heterogeneity in cortical progenitor division statuses, actual experimental data represent a mixture of these three cases. In Case 3, where neurogenic progenitors dominate, half of the clones are expected to consist of one or two neurons. Therefore, twice the proportion of one- or two-neuron clones serves as an estimate of the ratio of neurogenic progenitors at the time of retroviral injection. **B**. Representative coronal section showing three labeled clones (RFP in magenta, GFP in green) with spatial separation and no overlap. **C**. EdU labeling of postnatal (P1, P3, P6) mice and rats reveals co-labeling with Olig2 (glial progenitors) but not NeuN (neurons), suggesting no neurogenesis in the postnatal period in both species. **D**. Neuron-progenitor ratio defined by Pax6 (green) and beta III tubulin (magenta) expression demonstrates developmental stage correspondence between species with a two-day difference. E12.5 in mice and E14.5 in rats are the corresponding timing of onset of neurogenesis for the majority of cortical progenitors (yellow background). Scale bars: 1 mm (low magnification, **B**), 100 μm (magnified images, **B**), 50 μm (**C**), 200 μm (**D**).

**Figure S5.**
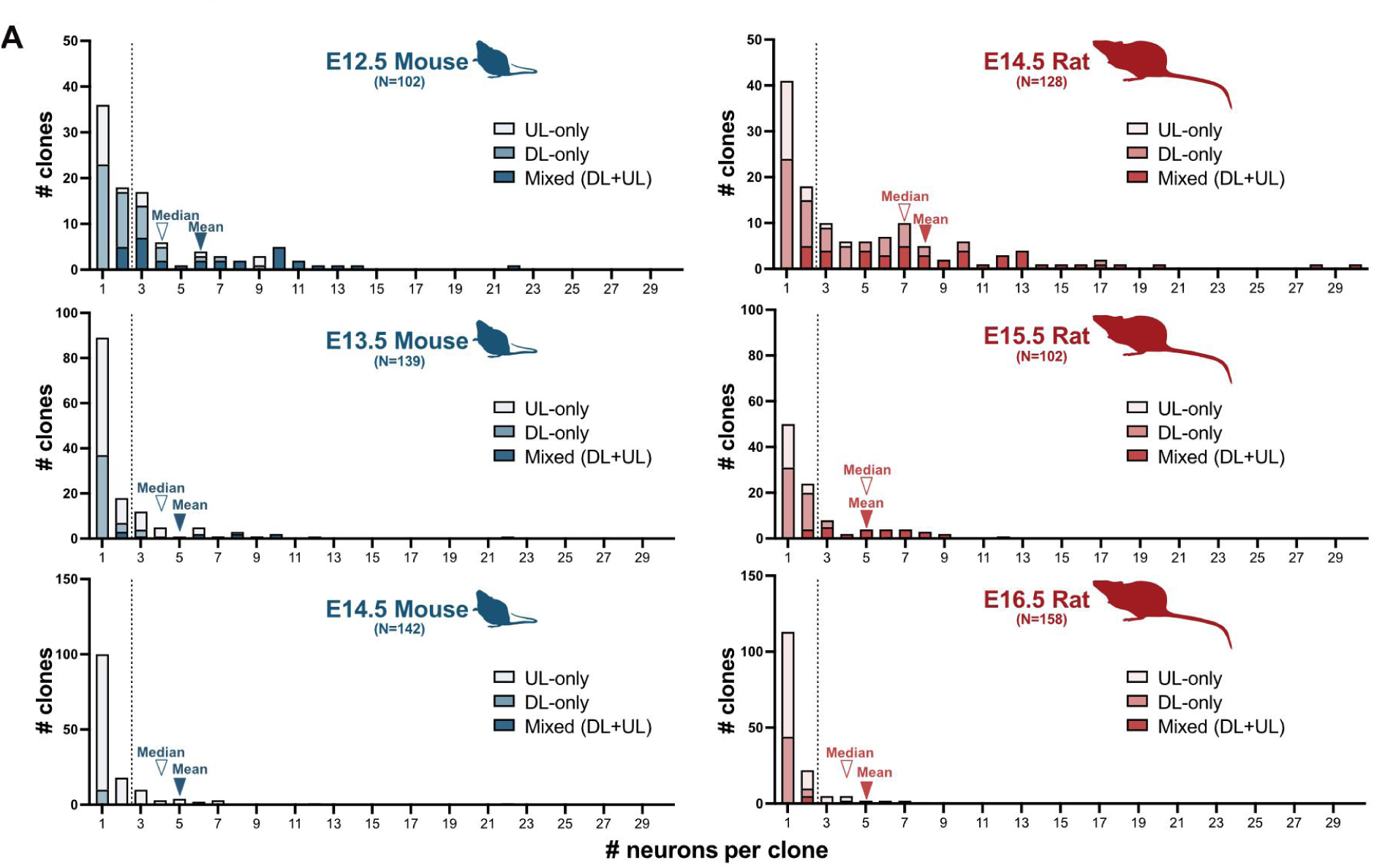
Clonal analysis with different retroviral injection timings, related to Figure 3. **A-F**. Distribution of clonal sizes in mice and rats with retroviral injections at different stages. Clone categories are visualized using color intensity.

**Figure S6.**
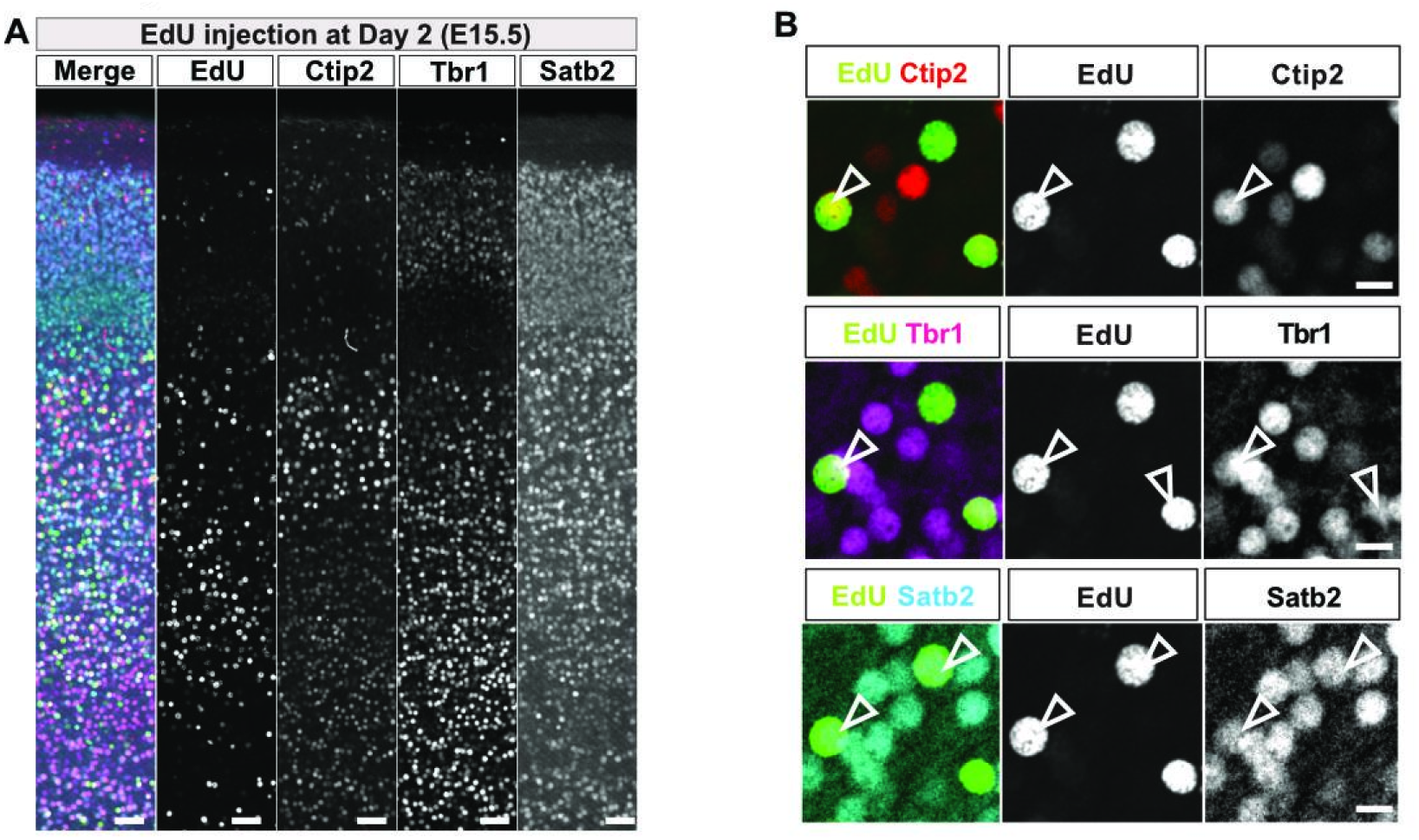
Marker immunoreactivility of EdU-labeled neurons, related to Figure 4. **A**, **B**. Representative images of cortical neurons co-labeled with EdU and subtype-specific markers. Open arrowheads indicate double-positive cells. Scale bars: 500 μm (**A**), 10 μm (**B**).

**Figure S7.**
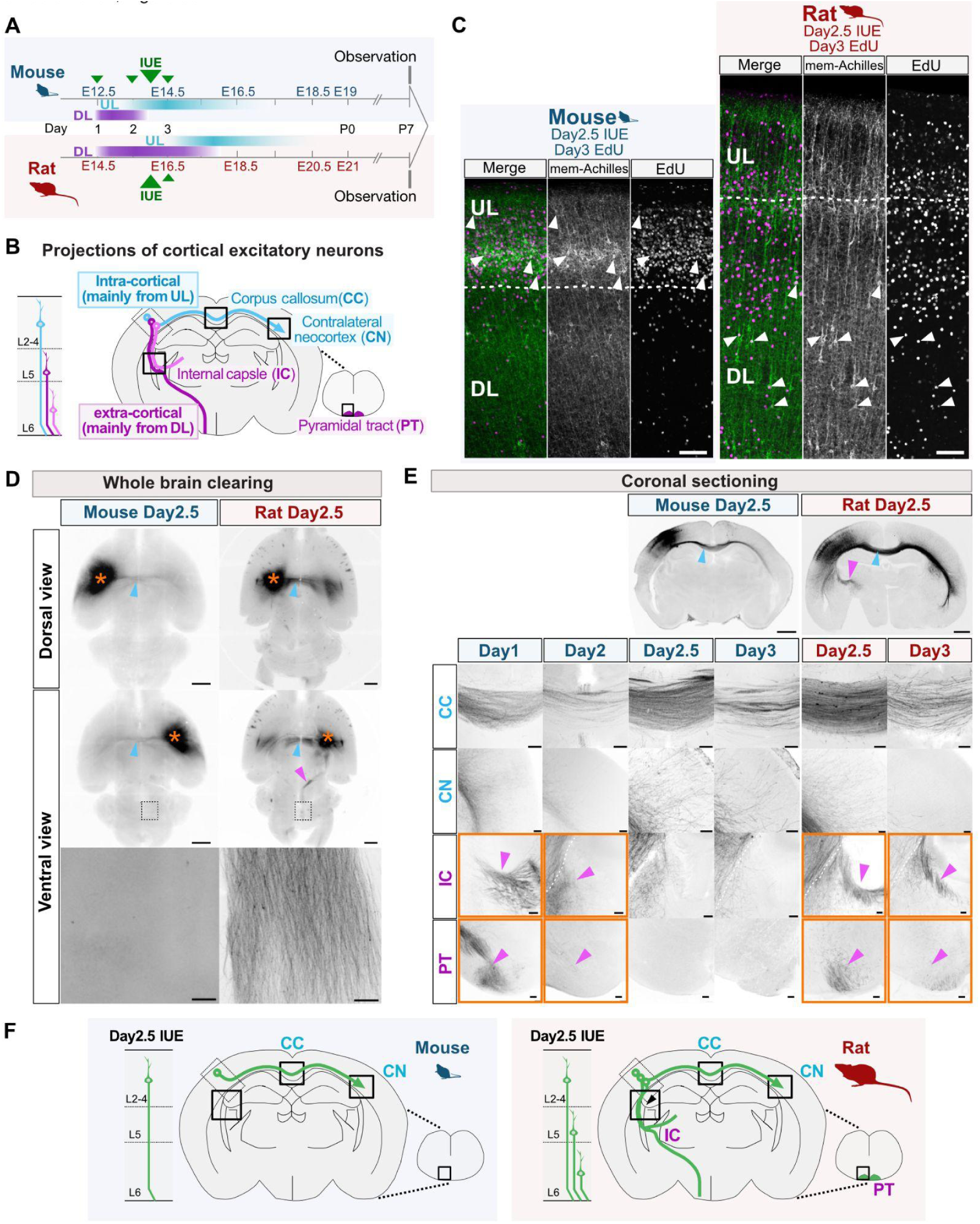
Projection identity of neurons generated at Day 2.5-3 of neurogenesis in mice and rats by *in utero* electroporation of a membrane anchored fluorescent protein, related to Figure 4. A. Schematic of IUE timing and sacrifice for axon projection analysis. **B**. Major projection targets of cortical excitatory neurons: corpus callosum (CC), contralateral neocortex (CN), internal capsule (IC), and pyramidal tract (PT). **C**. The neurons electroporated at Day 2.5 are co-labeled with EdU injected at Day 3 in both species. **D**. Tissue-cleared whole-brain images (dorsal and ventral views) show distinct projection patterns. In rats, prominent ipsilateral projections to the brainstem (magenta arrowheads) are unique. **E**. Coronal sections confirm unique descending projections in rats and their absence in mice after Day 3 IUE. **F**. Summary illustration of projection differences between mouse and rat neurons generated at Days 2.5–3.Scale bars: 1 mm (low magnification, **C** and **D**), 100 μm (magnified images, **D** and **E**).

**Figure S8.**
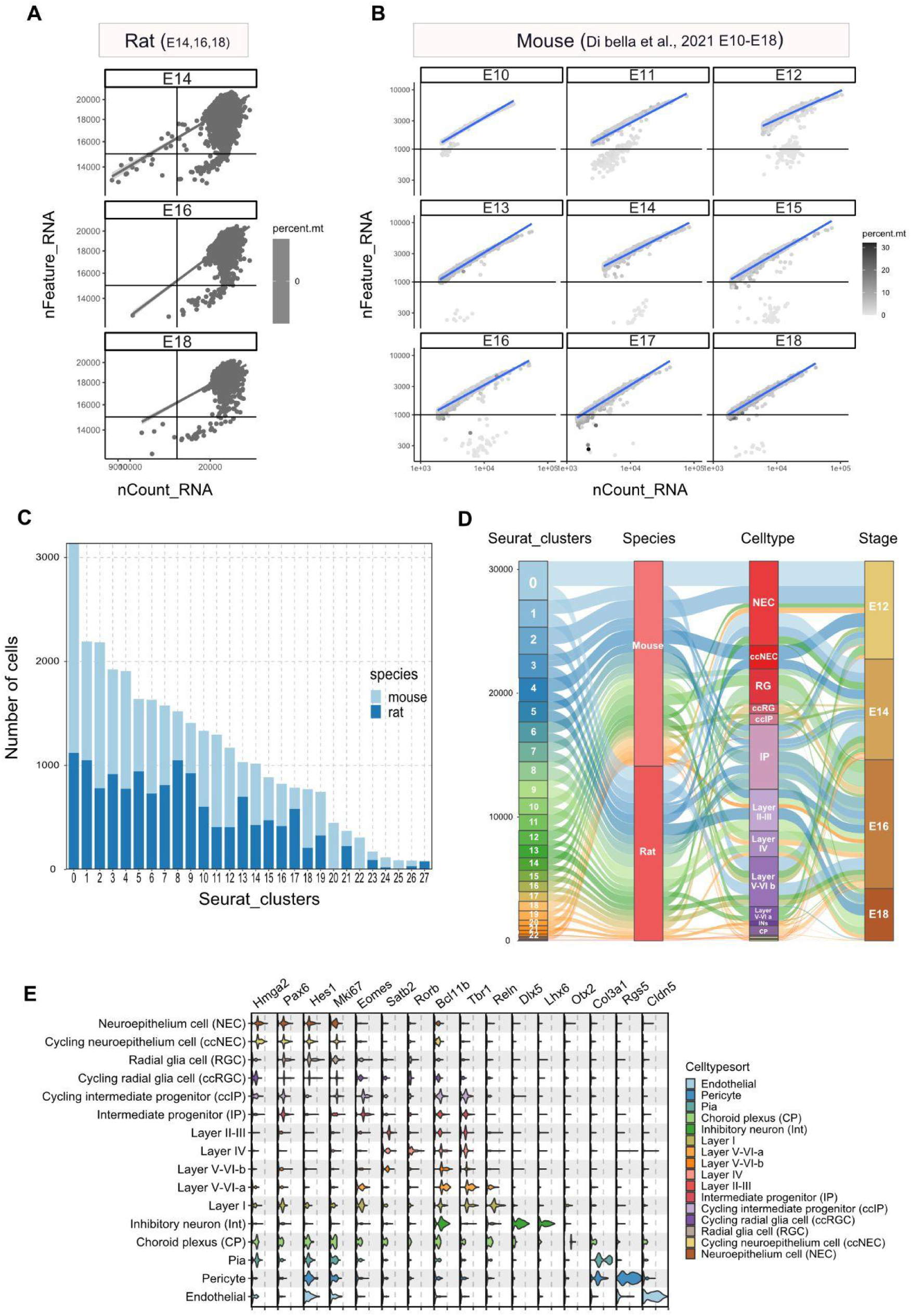
Single cell RNA sequencing of mouse and rat cortical cells, related to Figure 6. **A-B**. Distribution of detected genes, UMI counts, and mitochondrial read percentages in rat (**A**) and mouse (**B**) datasets. **C**. Cell numbers in UMAP clusters for both species. **D**. Relationship of clusters, species, cell types, and developmental stages. **E**. Expression of canonical markers in annotated cell types.

**Figure S9.**
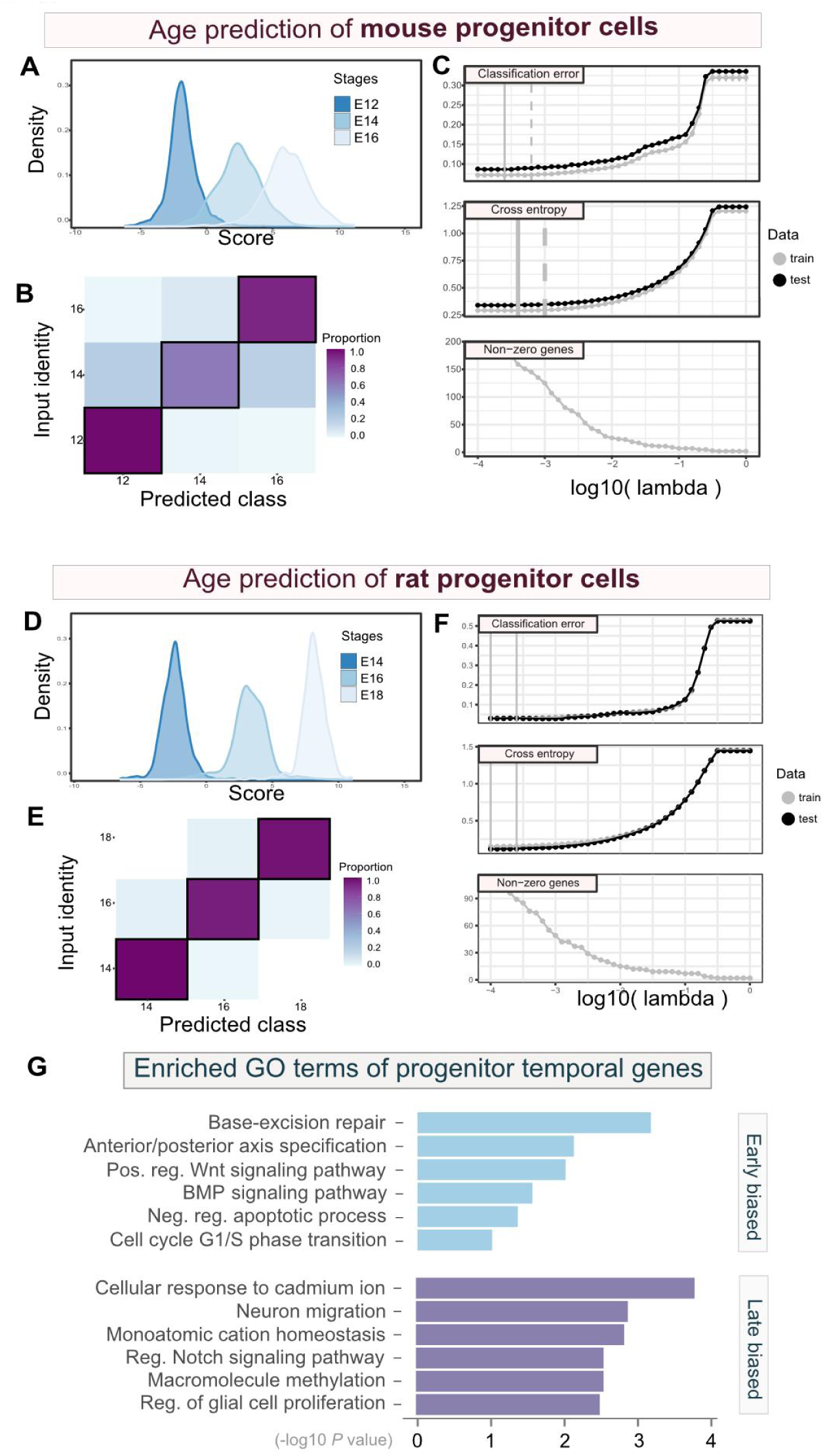
Identification of progenitor age genes using the ordinal regression model, related to Figure 6. **A-F**. Expression of progenitor age genes predicts sampling stages of progenitor cells in mice (**A-C**) and rats (**D-F**). **G**. Gene ontology analysis of progenitor temporal (age) genes.

**Figure S10.**
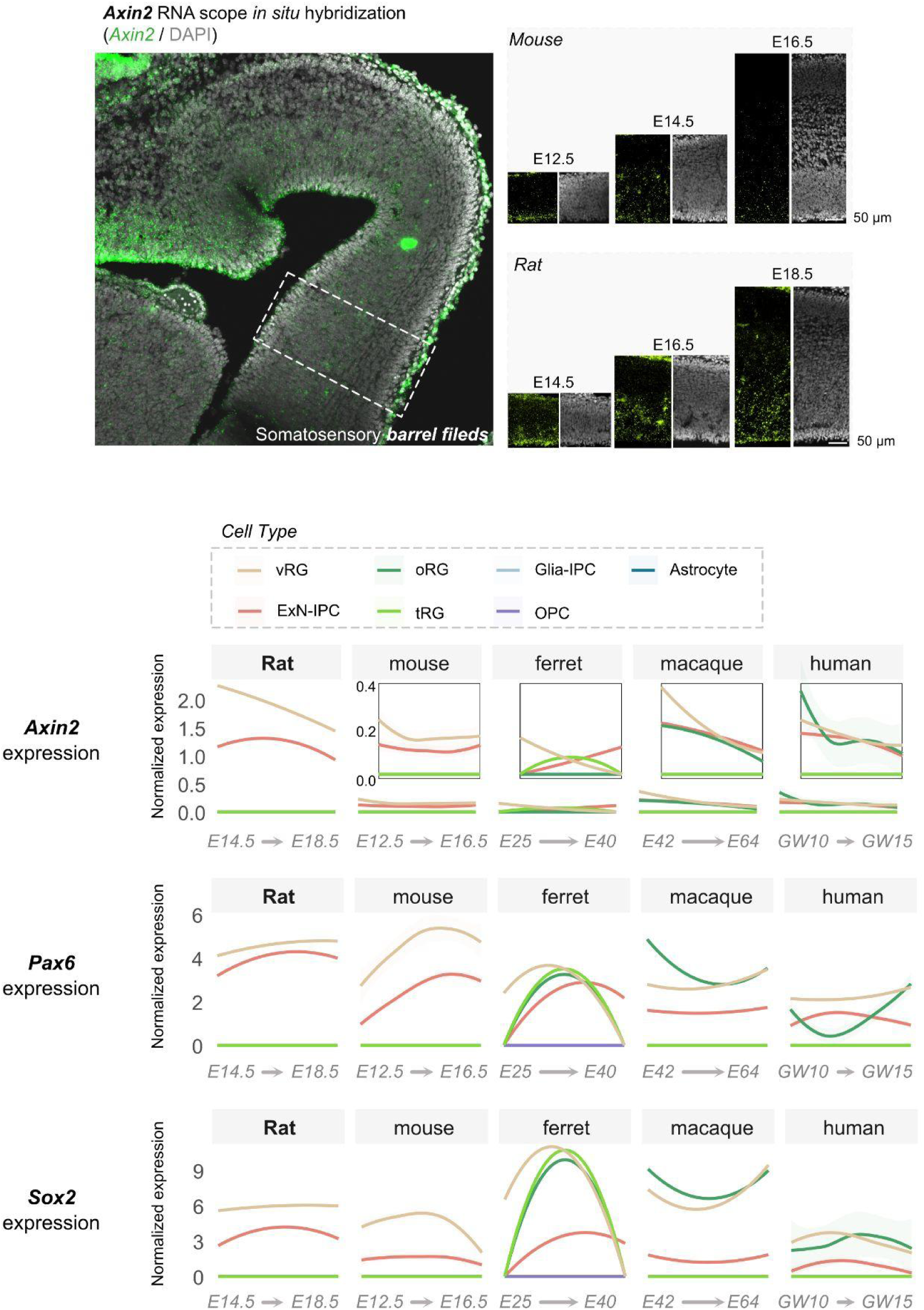
Significantly elevated expression of Axin2 in rat RG progenitors compared to mice and other mammals, related to Figure 7. RNAscope *in situ* hybridization was performed using an *Axin2* probe on coronal sections of the somatosensory cortex from mouse (E12.5, E14.5, E16.5) and rat (E14.5, E16.5, E18.5). Enlarged views of the barrel field (outlined by white dashed lines) are shown for each time point. Consistent with the analysis in Figure 7, single-cell RNA sequencing was conducted on rat and other mammalian species (using previously published datasets). Normalized expression values for Axin2, Pax6, and Sox2 across cell types are presented. Expression data were regressed out with sequencing depth and other potential technical variables as described in the Materials and Methods section.

**Figure S11.**
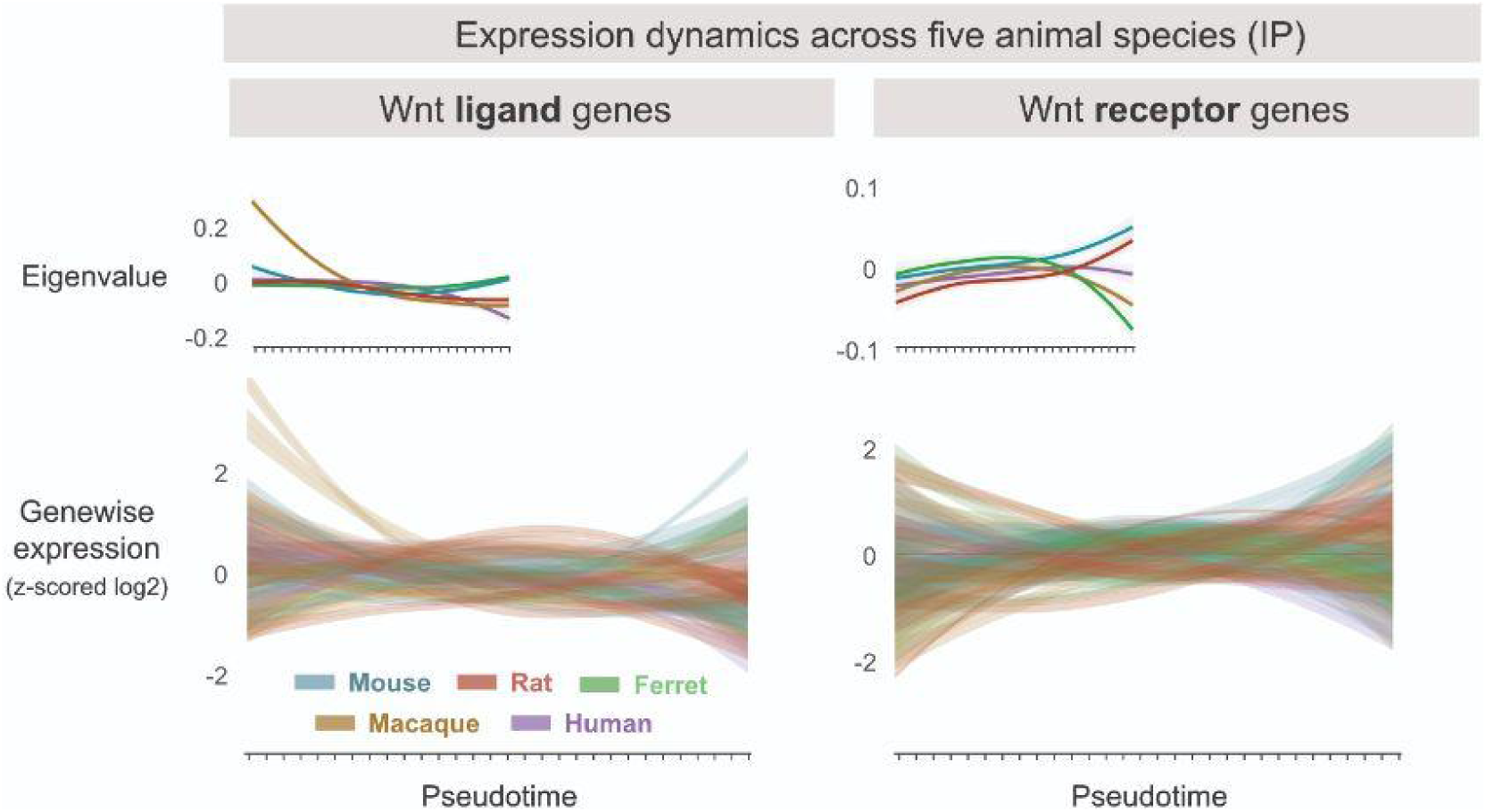
Conserved expression of Wnt-related genes in IP among all five mammalian species, related to Figure 7. Expression dynamics of Wnt ligand genes (n = 9) and Wnt receptor genes (n = 11) in excitatory neuron intermediate progenitor cells (IP) across five mammalian species. Gene expression values were log-normalized and z-scored across pseudotime to illustrate temporal dynamics, as described in the Methods section. Module eigenvalues, representing the first principal component of each gene expression matrix, are displayed in the upper left corner of each plot.

**Figure S12.**
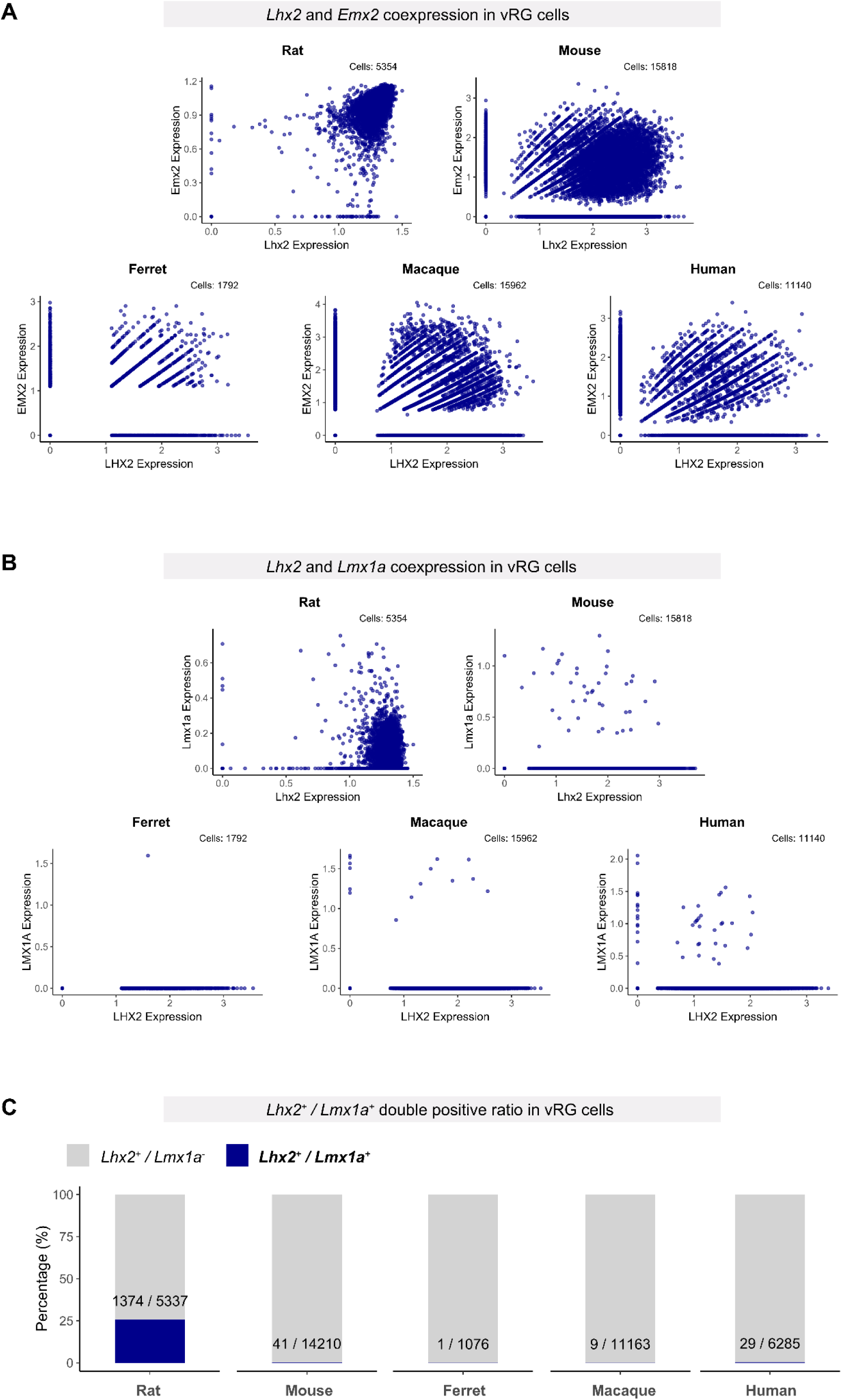
Gene coexpression patterns in ventricular radial glia (vRG)across mammalian species, related to Figure 7. **A.** Coexpression patterns of *Emx2* and *Lhx2* in vRG cells across five mammalian species (rat, mouse, ferret, macaque, and human). Expression values were retrieved from the SCT assay in each regressed object. **B.** Coexpression patterns of *Lmx1a* and *Lhx2* in vRG cells across five mammalian species (rat, mouse, ferret, macaque, and human). Expression values were retrieved from the SCT assay in each regressed object. **C.** Double-positive ratio of *Lhx2* and *Lmx1a* in vRG cells across five mammalian species. Cells with gene expression values above 0.1 for a given gene were defined as positive for that gene.

